# Combinatorial encoding of odors in the mosquito antennal lobe

**DOI:** 10.1101/2022.06.10.495686

**Authors:** Pranjul Singh, Shefali Goyal, Smith Gupta, Sanket Garg, Abhinav Tiwari, Varad Rajput, Alexander Shakeel Bates, Arjit Kant Gupta, Nitin Gupta

## Abstract

Among the cues that a mosquito uses to find a host for blood-feeding, the smell of the host plays an important role. Previous studies have shown that host odors contain hundreds of chemical odorants, which are detected by different receptors on the peripheral sensory organs of mosquitoes. But how individual odorants are encoded by downstream neurons in the mosquito brain is not known. We developed an *in vivo* preparation for patch-clamp electrophysiology to record from projection neurons and local neurons in the antennal lobe of *Aedes aegypti*. Combining intracellular recordings with dye-fills, morphological reconstructions, and immunohistochemistry, we identify different sub-classes of antennal lobe neurons and their putative interactions. Our recordings show that an odorant can activate multiple neurons innervating different glomeruli, and that the stimulus identity and its behavioral preference are represented in the population activity of the projection neurons. Our results provide the first detailed description of olfactory neurons in the central nervous system of mosquitoes and lay a foundation for understanding the neural basis of their olfactory behaviors.

## Introduction

*Aedes aegypti* has evolved as an anthropophilic mosquito species and a global vector of multiple diseases including dengue, zika fever, and chikungunya (McBride et al., 2014; Aubry et al., 2020). Along with temperature, humidity, and visual cues, olfaction plays a crucial role in the host-seeking behavior of mosquitoes (Rudolphs, 1922; McMeniman et al., 2014; Van Breugel et al., 2015; Liu and Vosshall, 2019; Vinauger et al., 2019). Volatile chemical molecules released from skin or present in breath guide mosquitoes toward their host (Cardé, 2015; Majeed et al., 2016; Raji et al., 2019; Zhao et al., 2022). Like other insects, mosquitoes also use olfaction for finding food, mates, and oviposition sites (Ponnusamy et al., 2008; Fawaz et al., 2014; Peach et al., 2019; Lahondère et al., 2020; Melo et al., 2020; Mozūraitis et al., 2020; Ye et al., 2021b).

The main olfactory organs in mosquitoes, like most insects, include the antennae and the maxillary palps (Bohbot et al., 2014; Raji et al., 2019). The axons of the sensory neurons project to the antennal lobe (AL) (Riabinina et al., 2016), where they synapse with broadly two classes of neurons: projection neurons (PNs) and local neurons (LNs) (Ignell et al., 2005). The insect AL, analogous to the olfactory bulb in vertebrates, is compartmentalized into glomeruli, such that each glomerulus receives input from a specific type of receptor neurons (Distler and Boeckh, 1997; Anton et al., 2003; Couto et al., 2005; Ghaninia et al., 2007a; Ye et al., 2021a). Although very little is known about the numbers and the morphologies of LNs and PNs in mosquitoes (Ignell et al., 2005), studies on other insects have shown that insect AL typically contains a few hundred LNs and PNs (Seki and Kanzaki, 2008; Chou et al., 2010; Singh et al., 2019; Bates et al., 2020; Schlegel et al., 2021). LNs innervate multiple glomeruli and provide lateral interactions within the AL (Chou et al., 2010; Mohamed et al., 2019; Fusca and Kloppenburg, 2021; Kymre et al., 2021). PNs innervate one or a few glomeruli and carry the output of the AL to the mushroom body and the lateral horn (Bates et al., 2020).

How are odors encoded by the PNs? Prior work in other insects suggests a combinatorial code, with each odor activating multiple PNs (Stopfer et al., 2003; Wilson et al., 2004), which emerges from combinatorial activity at the level of the receptors (Hallem and Carlson, 2006; Raman et al., 2010; Bisch-Knaden et al., 2018) along with lateral interactions within the AL. The vertebrate olfactory system also uses a combinatorial code for odors (Malnic et al., 1999). While a combinatorial code has obvious advantages for representing a large number of odors and for flexibly associating them with approach/avoidance behaviors (Masse et al., 2009; Choi et al., 2011; Warth Pérez Arias et al., 2020), how it preserves innate preferences to specific odors is less clear (Saha and Raman, 2015; Haverkamp et al., 2018). It may rely on stereotyped arborization of PNs in the higher brain centers, particularly the lateral horn, which is analogous to the mammalian cortical amygdala (Frechter et al., 2019; Bates et al., 2020). An alternative to a combinatorial code is a *labeled line*, in which a volatile molecule with a particular ecological role for a species may be detected by a specific receptor and further processed by a dedicated pathway in the brain, as observed for specific semiochemicals such as pheromones (Kurtovic et al., 2007; Ruta et al., 2010; Dweck et al., 2015; Taisz et al., 2022) or molecules indicating the presence of harmful species (Stensmyr et al., 2012; Huoviala et al., 2020).

It is unclear if the idea of labeled lines can be extended to olfactory behaviors that involve a large number of molecular components – for example, the attraction of a mosquito to a host. The host odor involves hundreds of different types of molecules (Cork and Park, 1996; Bernier et al., 1999, 2000; Gallagher et al., 2008; Dormont et al., 2013). Moreover, the types and the proportions of components vary considerably between different host species (Verhulst et al., 2018; Zhao et al., 2022) and among different members of the same host species (Syed and Leal, 2009; Obaldia et al., 2022). These components, even individually, can be detected by the mosquito olfactory system, as confirmed by the measurements of receptor responses (Davis and Sokolove, 1976; Ghaninia et al., 2007b, 2008; Lu et al., 2007; Hill et al., 2009; Syed and Leal, 2009; Carey et al., 2010; Tauxe et al., 2013; Majeed et al., 2016; Raji et al., 2019). The individual components can also generate specific behavioral preferences in mosquitoes (Smith et al., 1970; Geier et al., 1999; Bernier et al., 2007; Logan et al., 2008, 2010). It seems unlikely that each of these components has a dedicated neural pathway for itself in the mosquito brain, which appears similar to the fly brain in size (Ignell et al., 2005; Shankar and McMeniman, 2020). Previous studies of the mosquito AL have used either extracellular recording (Vinauger et al., 2018) or calcium imaging with a reporter in all cells or in the receptor neurons (Lahondère et al., 2020; Melo et al., 2020; Shankar et al., 2020; Zhao et al., 2022); with these approaches, it has not been possible to specifically examine the odor responses of PNs and LNs in mosquitoes.

Here we develop an *in vivo* preparation for patch-clamp recordings in mosquitoes to target PNs and LNs. Using post-hoc dye-filling we examine the glomerular identities and the detailed morphologies of the recorded neurons. We identify different sub-classes of PNs and LNs and compare their morphological and physiological properties. We analyze the odor responses of PNs and LNs and check the representation of the odor identity and the behavioral valence in the PN population. Our results provide a foundation for understanding the olfactory processing in the mosquito brain.

## Results

### Recordings from antennal lobe neurons

We developed an experimental preparation for *in vivo* recordings by immobilizing the mosquito inside a well in the middle of a custom-built recording chamber in such a way that the dorsal part of the mosquito head was accessible from the top of the well while the body of the mosquito remained below the surface (**Figure 1a**; see **Methods**). The perineural sheath from the dorsal and dorso-lateral region around the AL was removed gently, taking care to minimize the damage to the tissue. The well was perfused with physiological saline to keep the brain healthy during the experiment. The antennae and the maxillary palps were kept dry and accessible for odor stimulation. To minimize mechanical disturbances during odor delivery, the total flow rate of the final air stream reaching the animal was kept unchanged during the switch between clean air and odorized air (see **Methods**). This preparation allowed us to target individual cell-bodies of PNs and LNs and record their spontaneous activities and responses to a panel of odors.

**Figure 1:**
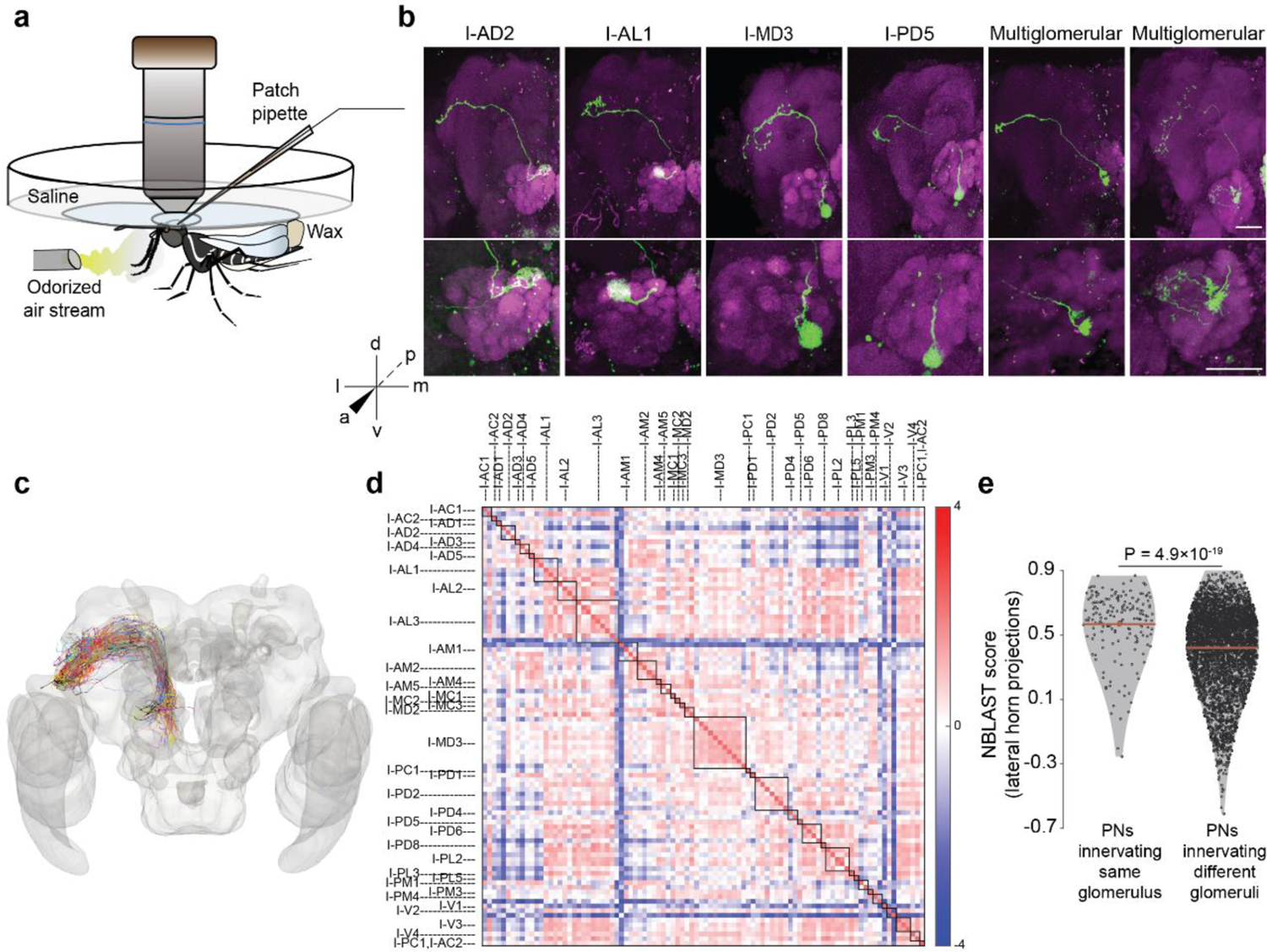
Morphological characterization of projection neurons (PNs) in A. aegypti. **a** Schematic showing the experimental preparation for *in vivo* patch-clamp recording. The animal was fixed with glue and wax to a recording chamber placed under the objective lens of an upright, fixed-stage microscope. Odors were delivered to the olfactory organs below the recording chamber and removed using a vacuum tube (not shown) placed behind the animal. Recordings were obtained from cell bodies present in the dorsal and the lateral regions around the antennal lobe (AL) using a glass pipette through a hole in the head cuticle. **b** Examples of some morphological sub-classes of PNs, including 4 uniglomerular, and 2 multiglomerular PNs. For each PN, the top image shows the complete morphology in a maximum-intensity projection of the image stack (the region displayed includes ipsilateral AL, mushroom body, and protocerebrum); the bottom image shows the dendritic innervations in a maximum-intensity projection from the AL. Co-ordinate axes: dorsal-ventral (d-v), anterior-posterior (a-p), and medial-lateral (m-l). Green: biocytin or lucifer yellow used to fill the cell; magenta: Dncad (neuropil marker). Scale bars, 50 µm. **c** Axonal reconstructions of 94 PNs registered to an *Aedes aegypti* brain template. Each PN sub-class is shown in a different color. **d** The matrix shows the NBLAST scores (after z-score normalization) indicating the morphological similarity in the protocerebral projections of pairs of PNs. PNs are ordered by glomerular identity, marked by the boxes along the diagonal and the labels along the axes. **e** NBLAST scores, indicating morphological similarity of projections to the lateral horn, were higher for pairs of PNs belonging to the same glomerulus (n = 171) than for pairs of PNs belonging to different glomeruli (n = 4200). Red lines: means, error bars: s.e.m.

### Morphological characterization of antennal lobe neurons

With our experimental preparation, we were able to target cell bodies in the anterodorsal and dorsolateral clusters around the AL. After the recording, we also attempted to fill dye in the recorded neuron followed by brain dissection and imaging for morphological identification. From a total of about 1250 mosquito preparations, we were successful in obtaining recordings along with morphological identification from 208 PNs and 53 LNs. We found that the distribution of cell bodies around the AL in *Aedes aegypti* is similar to *Drosophila melanogaster* (Wong et al., 2002; Das et al., 2008; Lai et al., 2008): the anterodorsal cluster contains cell bodies of PNs, while the dorsolateral cluster contains cell bodies of both PNs and LNs. By examining the dendritic innervations within the AL, we found that 201 of 208 recorded PNs were uniglomerular, with dense and complete innervation of the corresponding glomerulus (**Figure 1b**). Among the remaining multiglomerular PNs, we found five that innervated two glomeruli each, either entirely or partially (**Figure 1b**). We also found two PNs in the lateral cluster innervating more than two glomeruli (**Figure 1b**). Using the *Aedes* AL atlas provided by Ignell et al., 2005, we could assign glomerular identity to 175 uniglomerular PNs (see **Methods**). These PNs innervated 40 distinct glomeruli out of 50 described in the atlas.

Next, we checked the axonal projections of PNs in the higher brain centers. We were able to manually trace the axons of 94 PNs (93 uniglomerular covering 36 glomeruli and 1 multiglomerular); the others could not be traced due to low quality of dye fills or histology. The traces were registered to a female *Aedes* brain template (see **Methods**). Several antennocerebral tracts connecting the AL and protocerebrum have already been described in various insects (Mobbs, 1982; Homberg et al., 1988; Stocker et al., 1990). Among these, the inner antennocerebral tract (IACT), also called the medial antennal lobe tract (mALT), is known to be the most prominent tract. We found that the majority of the PNs sent their axons to the mushroom body calyx and the lateral horn through the IACT (**Figure 1c**). PN innervation at the lateral horn was in general more extensive than in the calyx. We calculated the pairwise neuronal similarity between the lateral horn projections of PNs (**Figure 1d**) using NBLAST and then compared the similarity between pairs of homotypic PNs (i.e., PNs innervating the same glomerulus, but not necessarily of the same morphology elsewhere) and pairs of heterotypic PNs. The NBLAST scores for homotypic PN pairs were moderately higher than for heterotypic PN pairs (0.57 ± 0.21 vs 0.42 ± 0.24, P = 4.9×10^-19^, rank sum test; **Figure 1e**), indicating that homotypic PNs have more similar projections in lateral horn.

All the identified LNs in our dataset projected ipsilaterally, although bilateral LNs are known in *D. melanogaster* (Chou et al., 2010; Schlegel et al., 2021). We analyzed the arborization patterns of 46 out of 53 LNs that had relatively clear stains and grouped them into four morphological sub-classes: *pan-glomerular* (covering entire AL; n = 11), *all-but-few* (covering almost entire AL except a few glomeruli; n = 26), *regional* (innervating a group of connected glomeruli; n = 7), and *patchy* (innervating a group of disconnected glomeruli; n = 2) (**Figure 2a**). The *pan-glomerular* and *all-but-few* sub-classes together can be considered equivalent to the “broad” class of LNs reported in *D. melanogaster* (Schlegel et al., 2021). In most cases of innervation of a glomerulus by an LN, the innervation covered the entire glomerulus, but in some cases, the innervation covered only a region within the glomerulus, suggesting compartmentalization within glomeruli. Next, we tried to check if some glomeruli are innervated more frequently or less frequently by LNs compared to other glomeruli. Given the difficulty in identifying glomeruli and in following weakly stained branches of LNs, we restricted this analysis to 14 landmark glomeruli (I-AD1, I-AD2, I-AM1, I-AM2, I-MD1, I-MD2, I-MD3, I-PC1, I-PD6, I-PL2, I-PL3, I-PL6, I-PM4, and I-V1) that were relatively easy to identify in different brains (**Figure 2b**) and 41 LNs whose branches were relatively clear. We found that, compared to other glomeruli, three anterodorsal glomeruli (I-AM1, I-AM2, and I-V1) have relatively fewer innervations by the LNs (**Figure 2c**).

**Figure 2:**
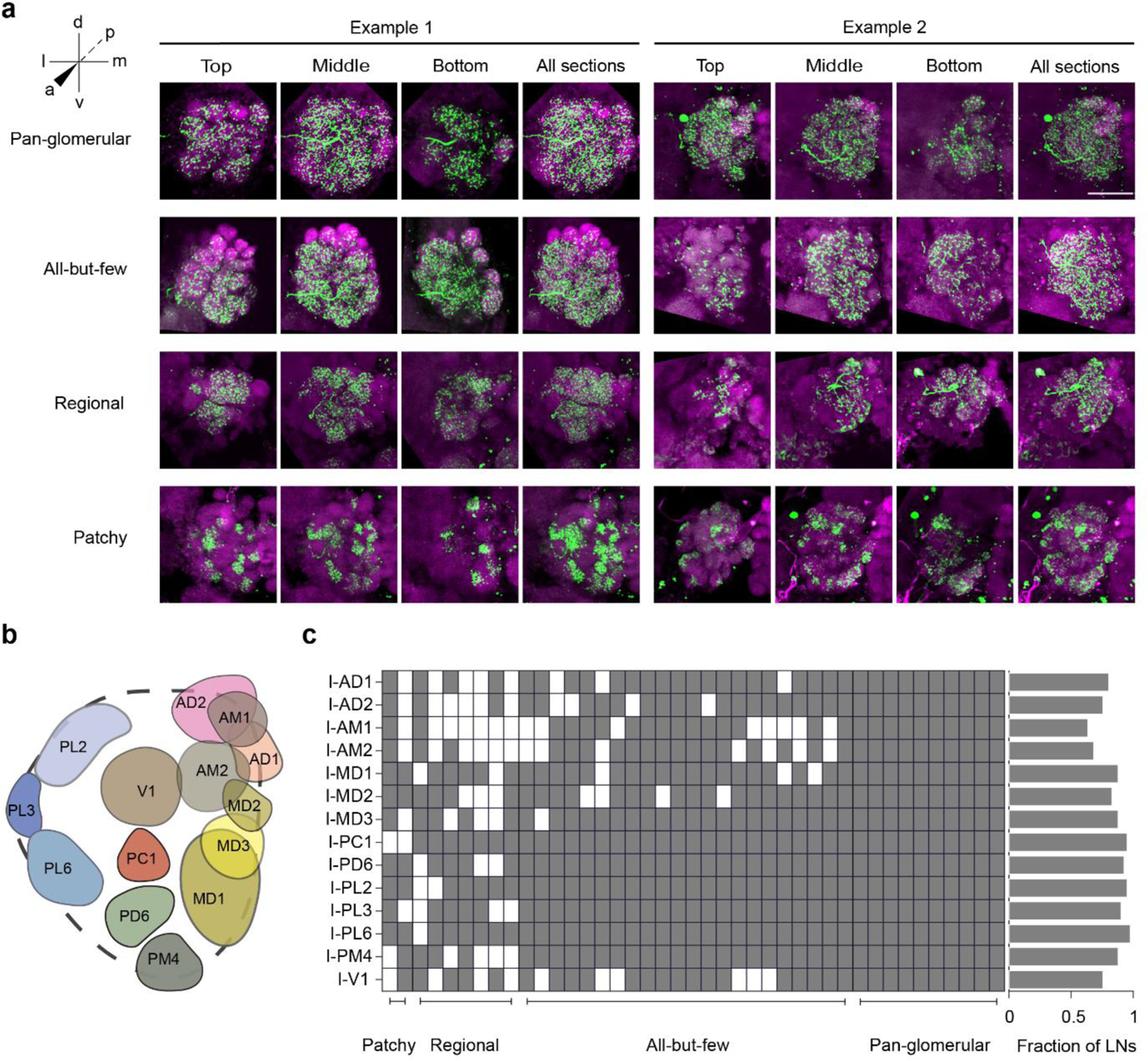
Morphological characterization of local neurons (LNs) in A. aegypti. **a** Each row shows two examples of LNs belonging to a different morphological sub-class. The first three images for each example show the maximum-intensity projections from different groups of planes from top to bottom, while the fourth image shows the projection from the entire AL. Green: biocytin/lucifer yellow (cell fill); magenta: Dncad (neuropil marker). Scale bar, 50 µm. **b** Schematic showing the positions of the 14 landmark (easily identifiable) glomeruli scattered across different regions of the AL. **c** Innervations of 41 LNs in the 14 landmark glomeruli (grey: innervation; white: no innervation). LNs are arranged according to the 4 morphological sub-classes. Right, each horizontal bar indicates the fraction of LNs innervating a glomerulus.

The tracer dye (biocytin) was injected only in the patched neurons. In some samples, however, we found that additional cell bodies, which were not targeted by the electrode, were labeled with the dye (**Supplementary Figure 1**). These sometimes included cell bodies in the ventral region of the AL, which was inaccessible in our preparation, and could not have been accidentally contacted by the electrode. Often the labeling of the additional cell bodies was accompanied by faint signals in multiple glomeruli, suggesting that the labeled neurons are PNs or LNs. Overall, this phenomenon was observed in at least 20 preparations, and points to the presence of gap junctions among the AL neurons in *Aedes aegypti*, as has been previously found in *D. melanogaster* (Huang et al., 2010).

### Electrophysiological classification of antennal lobe neurons

Taking advantage of the large dataset of identified PNs and LNs, we asked whether it may be possible to identify a recorded AL neuron as a PN or an LN, in the absence of a dye-fill, simply based on the electrophysiological recordings. We analyzed recordings obtained from morphologically identified PNs and LNs to test this idea. We classified spikes as isolated spikes or bursts (**Figure 3a**) and then extracted four electrophysiological features of isolated spikes: spike amplitude, spike half-width, after-hyperpolarization amplitude, and fraction of isolated spikes (**Figure 3b**; see **Methods**). These properties differed to different degrees between PNs and LNs (**Figure 3c-f**). Combining normalized values of the four properties into an electrophysiological feature vector for each neuron, we checked the correlations between pairs of neurons (**Figure 3g**). On average, we found positive correlations for pairs of PNs and for pairs of LNs, and negative correlations for PN-LN pairs (**Figure 3h**). LNs formed a more electrophysiologically homogenous group than PNs (within-group correlations of 0.59 ± 0.38 vs 0.23 ± 0.6, P = 4.5 × 10^-63^, rank-sum test). An unsupervised hierarchical clustering performed using the four properties was able to group the neurons into two broad classes matching the morphological classification of PNs and LNs with 95% accuracy (**Figure 3i**).

**Figure 3:**
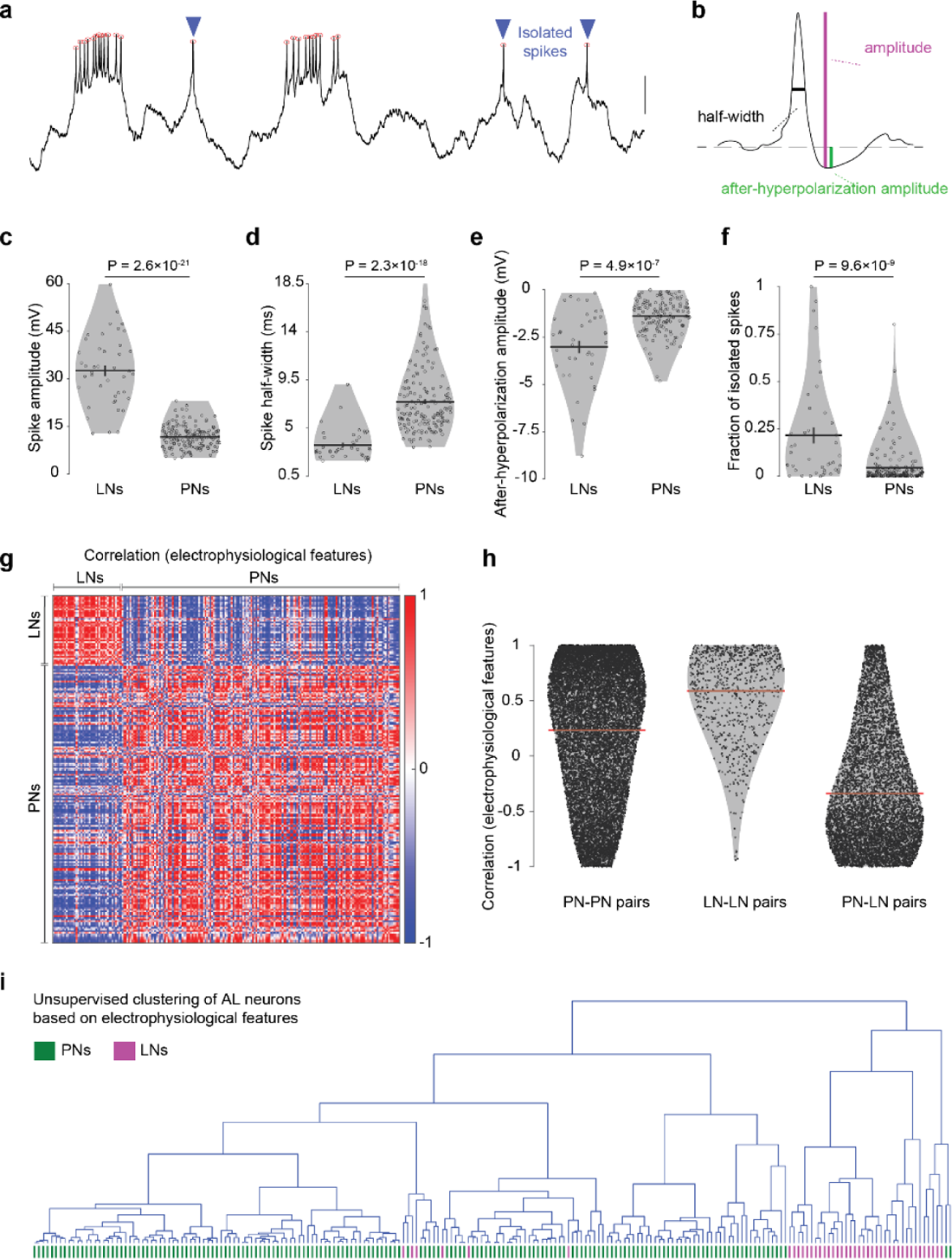
Electrophysiological classification of the antennal lobe (AL) neurons. **a** A representative recording trace (3.5s long) of an AL neuron. Red circles, detected spikes; blue arrowheads, isolated spikes (with no other spikes in ±200 ms); scale bar, 10 mV. **b** Illustration of electrophysiological features extracted from isolated spike. **c-f** LNs (n = 42) and PNs (n = 170) differed significantly in their values for the 4 electrophysiological features. P-values from rank-sum tests are displayed. **g** Correlations in the electrophysiological features between pairs of all AL cells, arranged according to the morphological class. **h** Correlation values were on average positive for PN-PN (n = 14365) and LN-LN (n = 861) pairs, and negative for PN-LN (n = 7140) pairs. **i** Unsupervised hierarchical clustering of AL neurons using the 4 electrophysiological features yields two broad classes corresponding to PNs and LNs with high accuracy; the actual identity of each neuron is indicated at the bottom in green (PN) or magenta (LN). Error bars: s.e.m.

Further, we checked if the electrophysiological properties depend on the morphological sub-classes within PNs and LNs. PNs belonging to the same glomeruli did not show higher correlations to each other than PNs belonging to different glomeruli (0.23 ± 0.60 vs 0.23 ± 0.61, P = 0.83, rank-sum test; **Supplementary Figure 2a**), thus showing that the electrophysiological properties do not depend on the glomerular identity of PNs. Within LNs (**Supplementary Figure 2b**), we found that neurons belonging to the *pan-glomerular* sub-class had significantly more similar electrophysiological properties than LNs belonging to different sub-classes (0.81 ± 0.18 vs 0.59 ± 0.38, P = 4.7 × 10^-4^, rank-sum test). Surprisingly, LNs from the *all-but-few* sub-class showed lower correlation to each other than LNs belonging to different sub-classes (0.49 ± 0.43 vs 0.59 ± 0.38, P = 0.02, rank-sum test), suggesting that the *all-but-few* sub-class is less homogenous than other sub-classes of LNs.

### Odor responses of projection neurons

PNs are the only neurons that carry olfactory information from the AL to higher brain areas. Therefore, it is important to understand how odors are represented by the PNs. We analyzed the responses of 175 morphologically identified uniglomerular PNs to a panel of 14 odors, including 8 components of human odor, 2 plant-derived volatiles, 1 carbon dioxide mimic, 1 oviposition attractant, 1 aggregation pheromone, and 1 synthetic repellent (see **Methods**), and solvents (mineral oil and water) (**Figure 4b**). In each PN, at least 6 trials were recorded for an odor, although not all odors could be tested with every PN. In a 10 second long trial, background activity was recorded for the first two seconds, followed by stimulation with odor for one second. Even in the absence of any stimuli, PNs showed spontaneous firing rates of different magnitudes (mean ± SD: 9.5 ± 9.3 Hz). Odors evoked a variety of responses in PNs, as represented by the examples in **Figure 4a**. The responses included bouts of excitation (increased firing) and bouts of inhibition (decreased firing). In many cases, the excitation was followed by inhibition (e.g., I-PD6 to DEET .1) or inhibition was followed by excitation (e.g., I-MD3 to 6MHO .01). In some cases, temporal patterns of multiple bouts of excitation and inhibition were observed (e.g., I-MD3 to DEET .1 or I-AL3 to PR ACID .01). Usually, the response started within 150 ms of the triggering of the odor delivery (this delay includes the time it took for the odor vapors to travel to the animal). However, some PNs responded to odors with much larger delays of 500-2000 ms. These delays cannot be attributed to a delay in odor delivery as we observed that the same odor that generated a delayed response in one PN could generate a fast response in another PN; the delays depended on specific PN-odor combinations (**Supplementary Figure 3**). In some specific cases, prolonged responses to odors were observed that lasted for several seconds after the termination of the odor stimulus (**Figure 4c**). Using a statistical criterion to determine if the odor response significantly different from the background firing (see **Methods**), we estimated the fraction of odors to which each PN responded (mean ± SD = 0.33 ± 0.25). These fractions differed for PNs belonging to different glomeruli, although most of them responded to multiple odors (**Figure 4d**).

**Figure 4:**
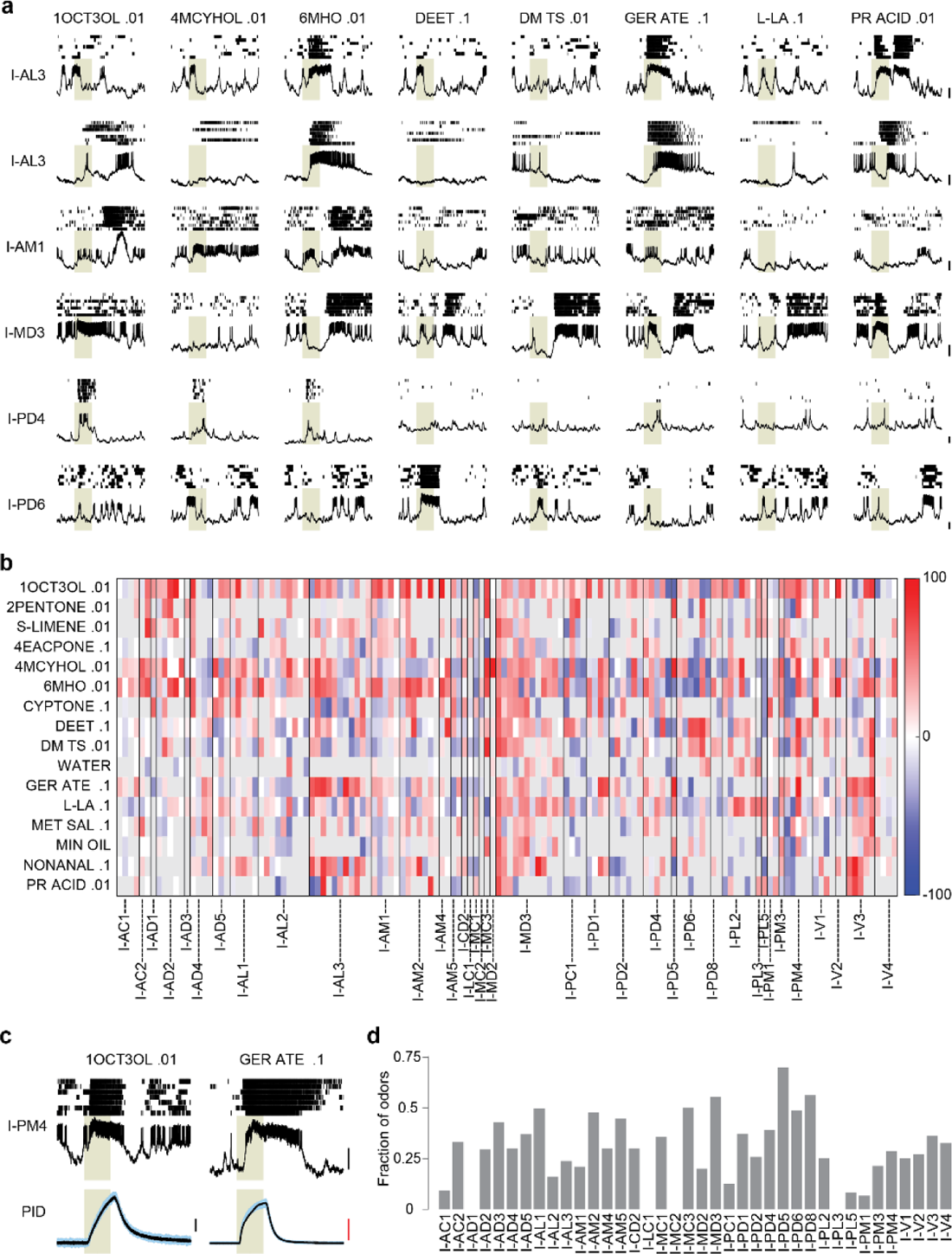
Odor responses of projection neurons (PNs) **a** Representative examples of odor responses from 6 PNs and 8 odors. Each panel shows a 5-s recording trace along and above it the raster of spikes from 7 trials. The first 2 PNs are homotypic (belonging to I-AL3 glomerulus), recorded from different animals, while the next 4 PNs belong to different glomeruli. Scale bar, 10 mV. Grey shaded region indicates the 1-s odor stimulation period. **b** Color-coded change in firing rate (spikes/s) evoked by 14 monomolecular odors and 2 solvents (mineral oil and water) in 138 PNs, each of which was tested with at least 5 odors. PNs are arranged by glomerular identity, separated by vertical lines. A grey value indicates that the odor was not tested in the PN. **c** Example of a PN showing a brief response to one odor and a prolonged response to another odor, which continues for 2-3 s beyond the odor stimulation. The bottom panel shows similar delivery profiles for both odors measured with a photo-ionization detector (PID). Scale bars, black: 10 mV; red: 100 mV. **d** The fraction of odors that elicited a response in a PN; values were averaged for PNs belonging to the same glomerulus.

Next, we evaluated the level of similarity between the odor responses of homotypic PNs. To ensure that the responses are compared using a uniform set of odors, we selected 6 odors that were most frequently tested in our dataset (6MHO .01, 1OCT3OL .01, DM TS .01, L-LA .1, DEET .1 and 4MCYHOL .01) and analyzed recordings from 64 PNs (covering 27 distinct glomeruli) in which 6 six odors were tested. We calculated the odor-evoked change in the firing rate for each of the 6 odors, combined these values to create a response vector for each PN, and calculated correlation coefficients between pairs of PN vectors (**Supplementary Figure 4a**). We found that the average Pearson correlation for PN pairs innervating the same glomerulus (0.26 ± 0.47) was higher than that of the pairs innervating different glomeruli (0.08 ± 0.47), indicating that odor responses of PNs within a glomerulus are on average more similar (P = 0.0016, rank-sum test; **Supplementary Figure 4b**). However, these correlation values had a large spread, indicating that the odor responses of PNs innervating the same glomerulus also varied in many cases. The variation could be because of the differences between differential inputs received by different PNs innervating the same glomerulus (indicating functional diversity among homotypic PNs in mosquitoes) or because of the differences between the individuals from which the PN recordings were obtained.

### Odor responses of local neurons

Compared to PNs, LNs showed lower spontaneous activity (2.6 ± 0.10 spikes/s; **Figure 5a**). LNs robustly responded to odors but these responses were in general of lower magnitude than observed in PNs (**Figure 5b**). Prolonged excitatory responses were rare in the LN population. In *D. melanogaster*, LNs have been reported to be broadly tuned (Wilson et al., 2004; Chou et al., 2010). Here, in A. *aegypti*, we found that LNs varied remarkably in the fraction of odors to which they responded (**Figure 5c**). In some cases, LNs showed reliable sub-threshold changes in the membrane potential, suggesting that LNs receive inputs for more odors than they respond to with spikes (**Figure 5d**). Next, we checked if responses were similar for LNs belonging to the same morphological sub-classes. We took a dataset of 22 LNs in which six common odors (same as those used for the PN analysis) were tested and estimated similarity as the correlation coefficient between pairs of LN response vectors. We found that odor responses of *pan-glomerular* LNs were significantly more similar to each other than responses of LNs belonging to different morphological sub-classes (0.55 ± 0.33 vs 0.0001 ± 0.52, P = 0.0001, rank-sum test; **Supplementary Figure 5a)**. Thus, *pan-glomerular* LNs are not only more homogenous electrophysiologically (**Supplementary Figure 2b**) but are also more homogenous in terms of odors responses compared to other morphological sub-classes, consistent with their broad innervation and higher probability of responding to an odor (**Figure 5c**).

**Figure 5:**
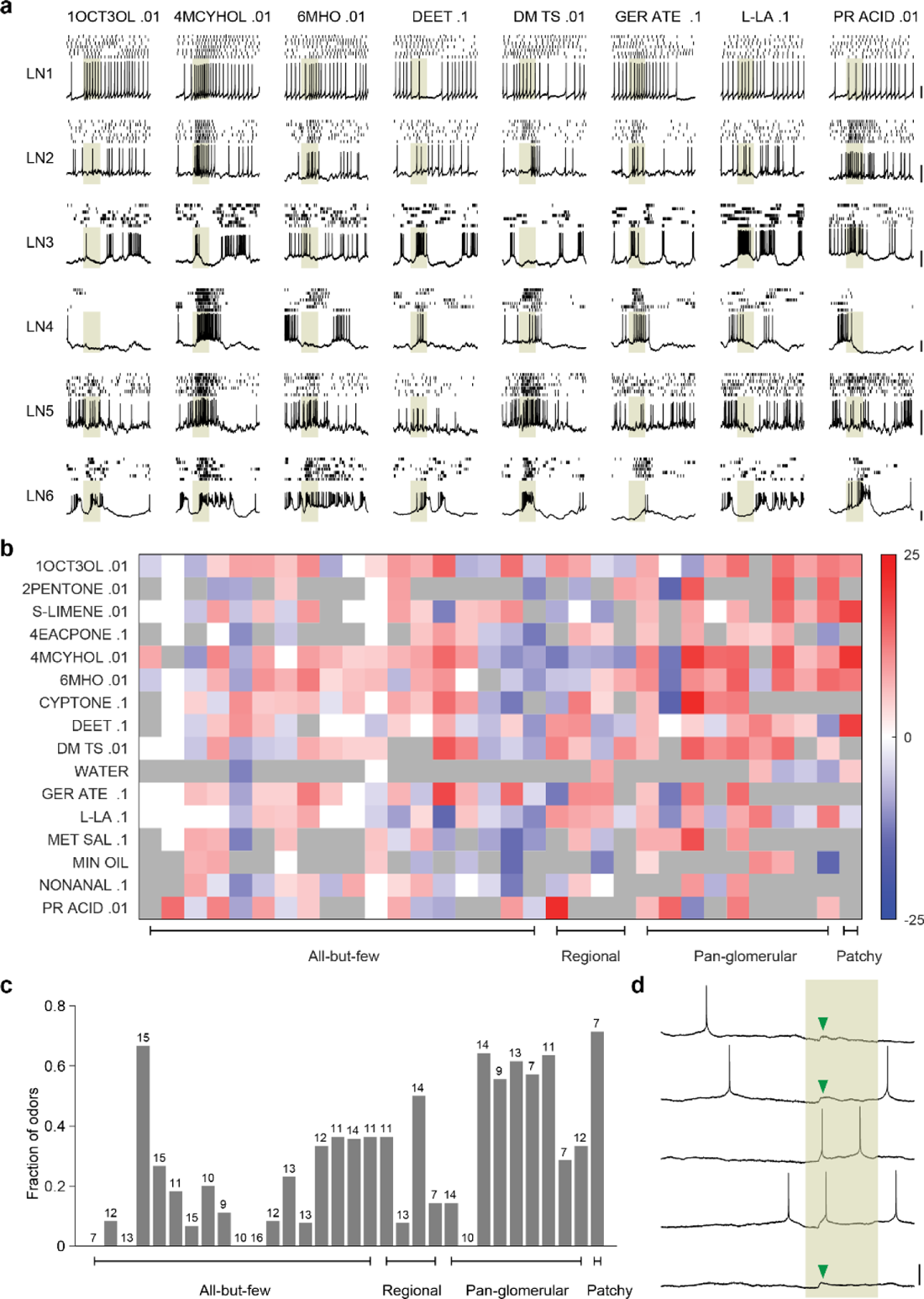
Odor responses of local neurons (LNs) **a** Representative examples of odor responses from 6 LNs and 8 odors. Each panel shows a 5-s recording trace along and above it the raster of spikes from 7 trials. Grey shaded region indicates the 1-s odor stimulation period. **b** Color-coded change in firing rate (spikes/s) of 32 LNs in response to 14 monomolecular odors and 2 solvents (mineral oil and water). LNs are arranged by morphological sub-classes. A grey value indicates that the odor was not tested in the LN. **c** The fraction of odors that elicited a response in each LN. The number on the bar indicates the number of odors tested for that LN. **d** Five trials of recording from an LN with 1-s stimulation of DM TS .01 (shaded region). The LN showed a spike soon after the odor onset in only 2 of the 5 trials, but showed an excitatory post-synaptic potential (arrowheads) around the same time in the remaining 3 trials. Scale bars, 20 mV.

As our patch recordings were performed in whole cell configuration, the cell body was often removed as the glass electrode was retracted after the recording. But in some cases, where we managed to cleanly detach the electrode from the cell body, we performed anti-GABA immunostaining to check if the recorded neuron was inhibitory. We found GABA-positive cell bodies in the lateral and ventral clusters around the AL (**Figure 6a**). A thick GABA-positive nerve fiber originating from the lateral group and going into the AL was also visible (**Figure 6a**). Among the recorded neurons with dye-fills and clear GABA stains, we had: 6 *pan-glomerular* LNs, of which 2 were GABA-positive and 4 were GABA-negative; 4 *all-but-few* LNs, of which 2 were GABA-positive and 2 were GABA-negative; and 8 PNs with cell bodies in the dorsal cluster, all of which were GABA-negative (**Figure 6b**). Presence of GABA-negative LNs suggests that they might be glutamatergic and inhibitory (Liu and Wilson, 2013) or cholinergic and excitatory (Shang et al., 2007).

**Figure 6:**
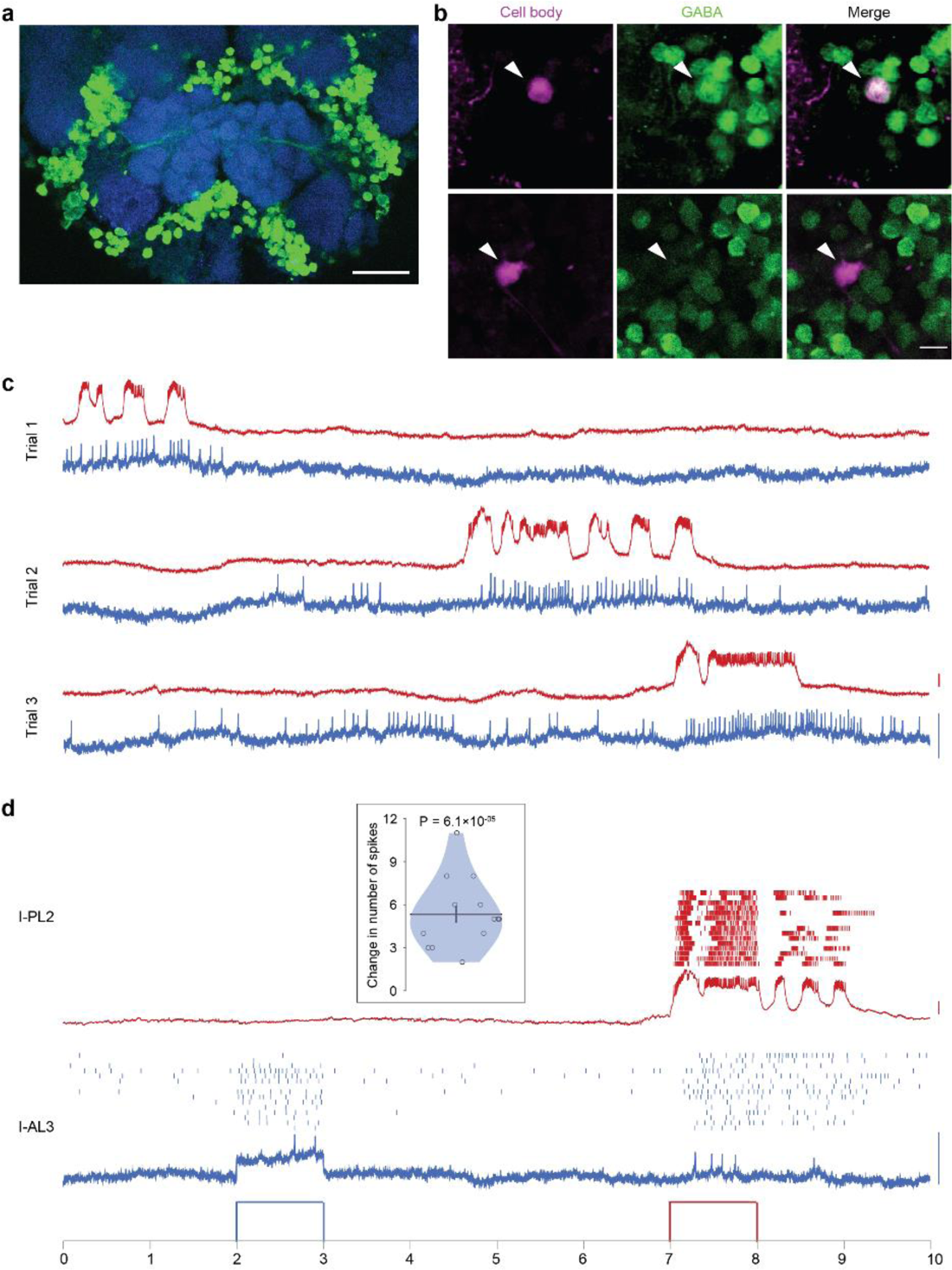
Lateral interactions in the antennal lobe (AL) **a** Image showing immunohistology with GABA antibody (green) and Dncad (blue, neuropil marker) around the AL in the *A. aegypti* brain. GABA-positive cell bodies can be seen in lateral and ventral clusters; a tract of GABA-positive fibers entering the AL from the lateral side can also be seen. Scale bar, 50 µm. **b** Optical sections from two brains (in two rows), in each of which an LN was recorded and filled (magenta) and immunostaining for GABA (green) was performed. One LN (top row) is GABA-positive and the other LN (bottom row) is GABA-negative. Scale bar, 10 µm **c** Spontaneous activity recorded simultaneously from an I-AL3 PN (blue) and an I-PL2 PN (red). Three trials of 10 s duration are shown. **d** Depolarizing current of 20 pA was injected during 2-3 s in I-AL3 (blue) and during 7-8 s in I-PL2 (red). Spike rasters for 15 trials (each 10s long) along with the recording trace of the first trial are shown for both neurons. Inset: the violin plot shows a significant increase in the number of spikes in I-AL3 during 7-8 s period when current is injected in I-PL2 (compared to 6-7 s period) over n = 15 trials. Scale bars, blue: 5 mV; red: 10 mV.

In one experimental preparation in which the antennae were severed to remove majority of the sensory input (palps were left intact), we succeeded in recording from one I-AL3 PN and one I-PL2 PN simultaneously (**Supplementary Figure 6b**). Although the spike patterns in the two neurons were not identical, there was a remarkable similarity in these two neurons (**Figure 6c**). To test whether this similarity was due to common inputs from other neurons or due to direct or indirect lateral interactions between the two neurons, we performed experiments in which we stimulated one neuron with current injection and observed the effect on the other. We found that while the injection of hyperpolarizing current in either cell had no effect on the other (**Supplementary Figure 6a**), stimulation of the I-PL2 PN with depolarizing current resulted in a reliable increase in the firing of I-AL3 PN (**Figure 6d**). This effect was directional, as the stimulation of the I-AL3 PN had no effect on the I-PL2 PN. The spike-triggered average of the I-AL3 PN’s membrane potential, triggered on I-PL2 PN spikes, suggested the lack of a direct connection between the two neurons (**Supplementary Figure 6c).** Together, these results point to an indirect lateral excitation from an I-PL2 PN to an I-AL3 PN, which may possibly be mediated by excitatory LN.

### Combinatorial coding of odors in the antennal lobe

The results so far indicate that a given PN or LN can be activated by multiple odors, and a given odor can activate multiple PNs and LNs (**Figure 4a, 5a**). The fraction of PNs activated by an odor depends on the odor identity: some odors like 6MHO .01, 1OCT3OL .01, and 4MCYHOL .01 activated about half of the PNs tests, while 4EACPONE .1 activated 16 % of cells, which was the smallest fraction among the odors tested (**Figure 7a**). Similarly, each odor activated multiple LNs with the exact number dependent on the odor-identity (**Figure 7a**). Interestingly, over the set of the odors tested, the fraction of PNs and the fraction of LNs activated by an odor were positively correlated (R = 0.63, P = 0.009, n = 16), suggesting that some odors in general elicited more widespread activity than others in the AL (**Figure 7b**).

**Figure 7:**
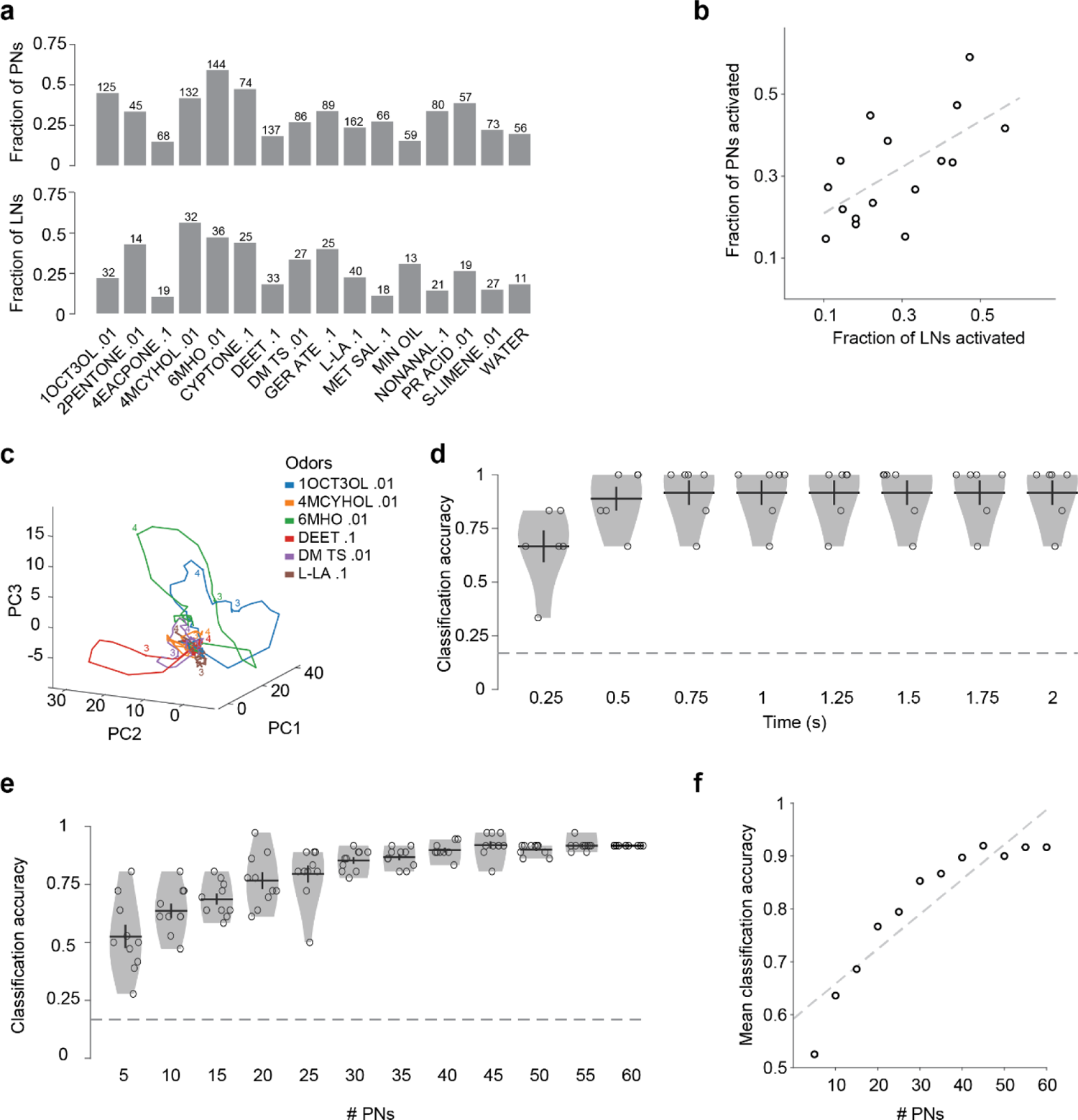
Combinatorial coding of odors in the PN activity. **a** Fractions of PNs (top) and LNs (bottom) activated by each odor. The number above each bar indicates the number of cells tested with that odor. **b** The fractions of PNs and LNs activated by odors were correlated (R = 0.63, P = 0.009, n = 16 odors). **c** Trajectories of PN responses to 6 odors projected on a 3-dimensional principal components (PC) space. The trajectories remained near origin before odor delivery (2 s timepoint) and began to diverge between 2 and 3 s timepoint, and gradually returned to origin (numbers on the plot indicate some timepoints). **d** Accuracy of odor classification based on PN population responses as a function of the response duration used. The accuracy for all durations exceeded the chance level (dashed line at 1/6). **e** The classification accuracy improves with an increase in the number of PNs taken for the analysis. **f** Correlation between mean classification accuracy (calculated in **(e)**) and number of PNs taken (R = 0.9, P = 2.7×10^-05^, n = 12).

The ability of the higher olfactory centers to distinguish between different odors depends on the inputs they receive from the PN population. We visualized the dynamic odor response of the PN population by reducing its dimensionality to 3 using principal component analysis (see **Methods**). The trajectories followed by different odors in the 3-dimensional space started at the same point but diverged over the course of the 1-s odor stimulation; the trajectories remained separated even after the end of the odor stimulus for another ∼1 s and gradually returned to the origin (**Figure 7c**). Thus, the odor identity seems be encoded by the temporally evolving responses of the PN population.

To confirm this, we checked how accurately the odor identity can be inferred from the PN population responses during individual trials. We considered the temporal response vectors (binned into 250 ms windows) of all PNs for increasing lengths of response periods following the odor onset, and then performed a classification analysis by comparing the population response during a given odor presentation to the templates constructed from other trials of the 6 odors in the dataset (see **Methods**). Compared to 1/6 classification accuracy expected by chance among 6 odors, we found nearly four times higher accuracy (0.67) within 250 ms of the response, which increased to 0.89 in 500 ms, and reached 0.92 in 750 ms, after which the inclusion of additional response periods did not change the performance (**Figure 7d**). These results suggest that the PN population encodes the identity of an odor within the first 500-750 ms of the odor response. To understand how many PNs are actually needed in the population for decoding the odor identity, we performed the same analysis by using different numbers of PNs chosen randomly. The results show that the odor identification among the 6 odors steadily improves as the number of PNs increases and begins to saturate around 40-50 PNs (**Figure 7e and f**). Thus, even though individual PNs respond promiscuously to odors, large groups of PNs can faithfully encode the odor identity in their temporally patterned responses.

### Innate valences of odors and their representation in the PN activity patterns

We next sought to understand the relationship between the PN population response and the behavioral preference elicited by an odor in *A. aegypti*. To evaluate innate preferences for the same odor concentrations that are used in electrophysiological recordings, we used a custom-made T-maze olfactometer designed to reveal both attractive and aversive preferences (see **Methods**) (**Figure 8a**). Briefly, it consisted of a large cuboidal chamber that was divided into two arms: one arm flushed with clean air and the other with odorized air stream. Mosquitoes were released into the center of the chamber and had to choose between the odorized arm and the control arm. A preference index (PI) for the odor was calculated as the number of mosquitoes in the odorized arm minus the number of mosquitoes in the control arm, divided by the total number of mosquitoes in the two arms. We found that 7 odors (4MCYHOL .01, CYPTONE .1, 2PENTNONE .01, 1OCT3OL .01, DM TS .01, 6MHO .01, and DEET .1) were significantly aversive (negative PI), one odor (L-LA .1) was significantly attractive (positive PI), and the remaining odors were behaviorally neutral at the tested concentrations (**Figure 8b**).

**Figure 8:**
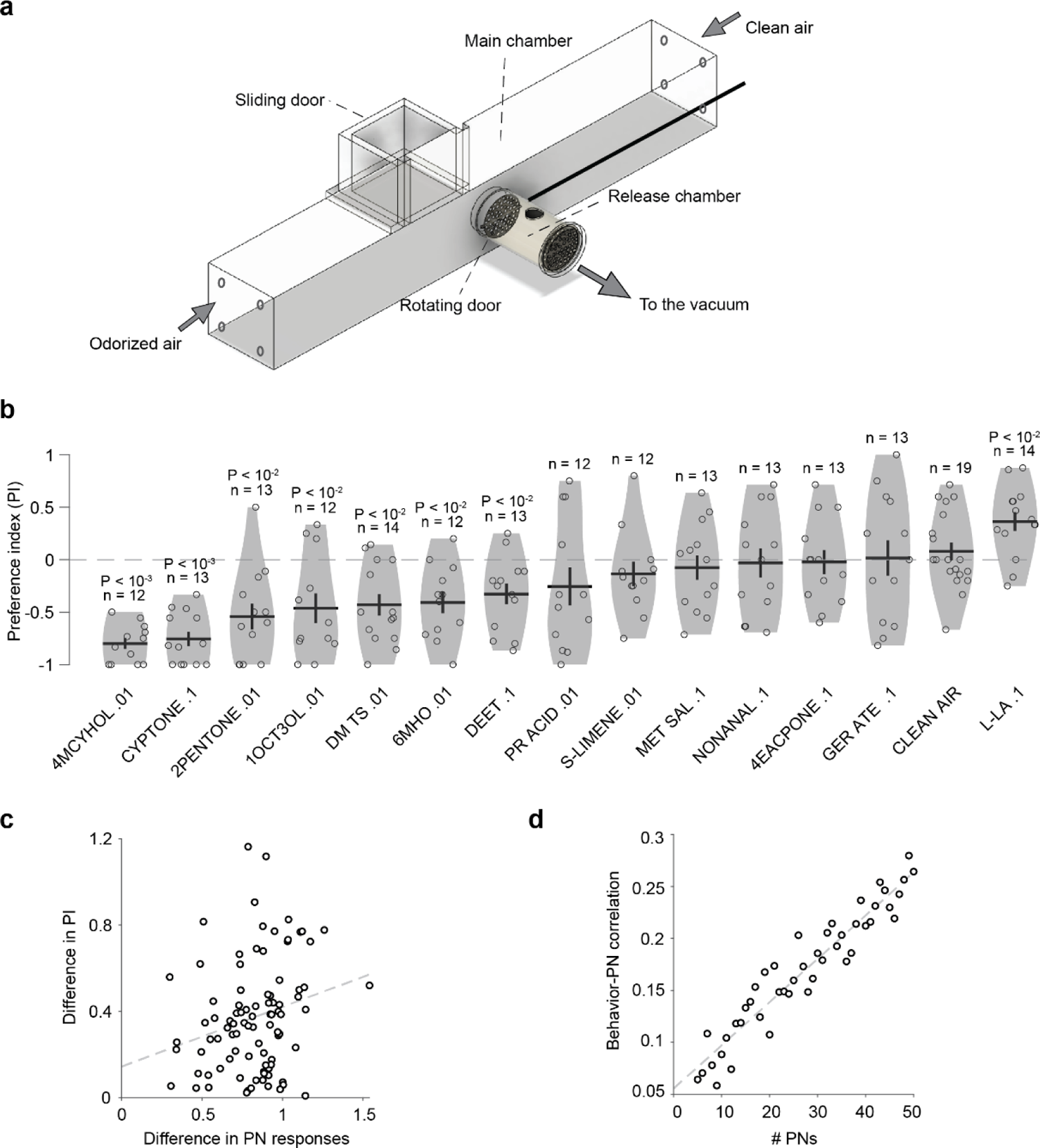
Representation of odor valence in the PN activity. **a** Behavioral chamber for evaluating the innate attractiveness and aversiveness of odors in *A. aegypti.* Female mosquitoes were placed in the release chamber 30 minutes before the experiment. At the start of the experiment, a rotating door separating the release chamber and main chamber was opened. One side of the main chamber was flushed with odorized air and the other side with clean air. After 5 minutes, the sliding double-door was closed, and the mosquitoes on each side were counted to calculate a preference index. **b** Preference index (PI) for 14 monomolecular odors. P-values are calculated from signed-rank test comparing PI to 0; insignificant P-values are not displayed. **c** Difference in preference indices (PI) is positively correlated with the difference in PN responses for odor pairs (R = 0.24, P = 0.02, n = 91). **d** The behavior-PN correlation increases with the number of PNs used in the analysis (R = 0.9, P = 3.2×10^-24^, n = 46).

Combining the behavioral and the electrophysiological data, we checked if the odors generating similar behavioral preferences also generated similar activity in PNs. For each pair of odors, we calculated the difference in their PIs and the difference in their PN responses (calculated as 1 minus the correlation between the vectors of odor-evoked responses in PNs that were tested with both odors; see **Methods**). Over all pairs of odors, the difference in PIs was positively correlated with the difference in PN responses (R = 0.24, p = 0.023; n = 91 odor pairs from 14 odors; **Figure 8c**). Next, we asked how this behavior-PN correlation depends on the PN population size. To check this, while calculating the difference in PNs responses for a pair of odors, instead of using all the PNs that were tested with the two odors, we randomly subsampled a given number of common PNs, and recalculated the behavior-PN correlations from this reduced PN population; this sampling was performed 20 times to estimate the average behavior-PN correlation for each number of PNs. The results showed that the average behavior-PN correlation increases with the number of PNs considered (R = 0.95, p = 3.2 x 10^-24^; **Figure 8d**).

## Discussion

We recorded the responses of LNs and PNs to several odors and also measured the behavioral preferences for these odors at the same concentrations. The behavioral assays were performed using a custom T-maze olfactometer with a small stem (a small cylindrical release chamber). A long stem can be problematic with aversive odors as the animals might not come out of the release chamber to fly upwind and reach the decision point between the odor arm and the control arm. In our olfactometer, a small release chamber resulted in mosquitoes entering the decision-making zone more frequently. We used active air flow to precisely control the concentration of odor within the chamber. Our large dataset of neuronal responses, along with behavioral data, allowed us to explore the general principles of odor representation in the AL. We found that although an individual PN typically responds to multiple odors, the pattern of activity in the PN population is unique for each odor. These patterns for different odors are separated well enough to allow reliable odor discrimination. We also found that the odors become more discriminable when responses from more PNs are considered. Odor pairs that elicit similar activity in the PN population elicit similar behavioral preferences in mosquitoes. Together, these results support a combinatorial code for odors in the PN population.

We obtained the recordings by adapting the technique of *in-vivo* whole cell patch clamp recordings for the mosquito AL. Multi-electrode recordings and calcium imaging provide population level information. However, properties of individual neurons in a circuit and network interactions are difficult to assess with these techniques. Intracellular electrophysiology, on the other hand, provides better resolution about how different neurons in a circuit interact and data can be pooled to get a sense of the final population level output, especially in a relatively numerically simple nervous system such as that of an insect. Another advantage of using this technique is the ability to perform dye-fills and post-hoc histology that reveals the morphology and therefore the identity of the recorded neuron. This is particularly useful in systems where genetic labelling is not easy, such as with mosquitoes.

We found that the morphological organization of AL neurons in mosquitoes is similar to that of *Drosophila*, but with some differences. Unlike in *Drosophila* where more than half of the PNs are multiglomerular (Bates et al., 2020), 201 of the 208 PNs in *A. aegypti* we observed were uniglomerular; this may be related to the fact that cell bodies in ventral AL were not accessible in our preparation. We found many LNs with broad innervations (*pan-glomerular* and *all-but-few* sub-classes) and some LNs with selective innervations in glomeruli (*regional* and *patchy* sub-classes). Similar sub-classes have been described in *Drosophila* (Chou et al., 2010; Schlegel et al., 2021), with one notable difference: we did not observe any bilateral LN in our sampling from mosquitoes. Taking advantage of the detailed information available in intracellular recordings, we were able to extract four features of spikes that could predict whether a recorded neuron is a PN or an LN. This result is consistent with studies on other insects showing differences between PNs and LNs (Wilson et al., 2004; Kuebler et al., 2011; Lei et al., 2011; Meyer et al., 2013), and will help in identifying neurons in cases of unsuccessful dye-filling or damage to the brain tissue while handling.

Many of the odors used in this study have been at the center of mosquito olfaction research. For example, 6-methyl-5-hepten-2-one (6MHO, also known as sulcatone) is an important skin component and evolution of *A. aegypti* to bite humans has been attributed to increased expression of receptors for 6MHO (McBride et al., 2014). However, a high concentration of sulcatone in human odor has been reported to decrease attraction (Logan et al., 2008). In our behavioral experiments, 6MHO .01 was aversive and in our patch-recordings, we found that 6MHO .01 activates a large subset of PNs (**Figure 3d**). DEET is another odor that has been of great interest as the most widely used repellent against mosquitoes. Despite being in use for a very long time, the mode of action of DEET has been puzzling and appears to be species-specific (Leal, 2014; Afify and Potter, 2020). Using calcium imaging, Lahondère et al., 2020 recently showed that DEET activates AM2 glomerulus in *A. aegypti;* however, since the calcium reporter used in this study was driven by the ubiquitin promoter, it was not possible to pin down this activity to a specific cell-type within the glomerulus. With our recordings, we found that PNs innervating I-AM2, I-AM4, I-MD3, I-PD6 and I-PM4 respond to DEET (**Figure 3b**). In our behavioral experiments, mosquitoes were repelled by DEET vapors mixed with the background air stream. Together, our results show that DEET is detected as an odor in *A. aegypti* and a combinatorial activity in the PN population may be responsible for its aversive nature. The PN-odorant combinations tested in our study, along with the information about the cell-types and putative interactions in the mosquito AL, provide a foundation for understanding how a large number of behaviorally relevant odorant are processed in the mosquito brain.

## Methods

### Animal stock

All experiments were performed with *Aedes aegypti* (Linnaeus) Liverpool strain. Larvae were reared at 28±1° C and 60±10% relative humidity in 500 ml round plastic containers containing deionized water and fed on powdered fish food (TetraBits). Pupal cups containing 150-200 pupae were placed in a mesh cage (L:30 X B:30 X H:30 cm) for adults to emerge. Adults were fed on a 10% sucrose solution using a cotton feeder and kept at 25±5° C and 60±15% relative humidity under a photoperiod of 14 hr:10 hr (light:dark). Females were blood-fed using mice (*Mus musculus BALB/c*) and eggs were collected in plastic containers lined with filter paper strips. For experiments, non-blood-fed mated female mosquitoes of age 4-8 days were used.

### Animal preparation and electrophysiology

*In-vivo* whole-cell patch-clamp recordings were obtained from cell bodies around the AL in *Aedes aegypti*. The mosquito was anesthetized on ice and the legs were immobilized using wax. The animal was then transferred to an aluminum foil covering a small hole in a plastic cup. The foil above the hole was cut and adjusted to hold the mosquito in such a way that the head and the thorax of the mosquito were accessible from the top while the remaining part was hanging below. Wings and the proboscis were immobilized using wax and the proximal segment of the antennae was pasted to the anterior part of the head using an epoxy adhesive (Araldite). A piece of a thin plastic wrap with a small rectangular cut in the center was pasted on the mosquito above the head and thorax such that the dorsal part of the head remained exposed. The animal was then transferred from the plastic cup to a hole in a recording chamber, made of a petridish, and the sides of the plastic wrap were sealed onto the surface of the chamber. This way the dorsal part of the head was accessible from the top while the olfactory organs were below the plastic sheet. This ensured that the olfactory organs remained dry and exposed to odor stimulation during experiments. The well was then filled with saline and a window was cut in the exposed part of the head using a sharp tungsten needle. The tracheae were removed, and the perineural sheath was partially removed to expose the cell bodies. In most of the samples, a brief exposure to 0.5% collagenase type IV was used to facilitate de-sheathing (this did not cause any noticeable difference in the recorded activity of the cells). The brain was perfused continuously with saline bubbled with 95% O_2_/5% CO_2_. The saline composition was as follows (in mM): TES (12.5), Glucose (8), Sucrose (2), NaCl (120.5), Trehalose dihydrate (6.25), NaHCO_3_ (17), NaH_2_PO_4_ monohydrate (0.6). The pH was adjusted to 7.3 and osmolarity was set to 300-305 mOsm. The animal preparation was placed under Nikon FN-1 microscope with 40X water immersion objective. Patch-clamp electrodes (6.5-9 MΩ) were pulled from glass capillaries (1.5×0.86 mm) using electrode puller (Sutter Instrument P1000). Recording electrodes were filled with an internal solution composed of the following (in mM): Potassium aspartate (140), HEPES (10), MgATP (4), Na_3_GTP (0.5), EGTA (1.1), KCl (1), KOH (1). The pH of the internal solution was adjusted to 7.1 and osmolarity was set to 295 mOsm. Biocytin (0.5%) or Lucifer Yellow (0.1%) was added to the internal solution.

Recordings were acquired in current-clamp mode using a Multiclamp 700B amplifier (Molecular Devices), sampled at 20 kHz and low pass filtered at 10 kHz. In most cells, a small constant current was injected after breaking in to bring the resting membrane potential between −45 to −60 mV. Within an experiment, 7 trials were recorded for each odor. The total duration of one trial was usually 10 s and odors were delivered for 1 s after 2 s of trial onset. However, if a cell showed prolonged odor response, the duration of the trial was increased to make sure that the firing rate returns to baseline before the onset of the next pulse of odor. Tracer dye (0.5 % Biocytin or 0.1% Lucifer Yellow) was iontophoretically injected into the cells from which stable recordings were obtained. To fill the dye, a hyperpolarizing pulse (500 ms) of 3 nA was applied at 1Hz for 20-40 mins after recording. In paired recordings, one cell was filled with biocytin, and the other cell was filled with lucifer yellow.

### Histology

After the recordings, the brains were dissected out and fixed in 4% PFA at room temperature for 30-60 mins. Quick washes with PBS were followed by incubation in PBS containing 0.2% Triton-X (PBS-T) for 1 hour. Then, the brains were incubated in 5% normal goat serum at room temperature for 2-3 hr or kept at 4 ° C for overnight incubation. After this, the samples were incubated in 1:30 rat anti-DN-cadherin (DN-EX #8, Developmental Studies Hybridoma Bank) and 1:500 rabbit anti-GABA (A2052, Sigma-Aldrich) or 1:200 rabbit anti-Lucifer yellow (A5750, Molecular Probes) for 1-2 days at 4° C. Brains were washed in PBS-T for several hours at room temperature and incubated with 1:500 goat anti-rat with Alexa 405 (ab175671, Abcam), 1:500 goat anti-rabbit with Alexa 633 (A21070, Molecular Probes) or goat anti-rabbit with Alexa 488 (A11008, Molecular Probes) and 1:10^5^ Streptavidin with Alexa 488 (S11223, Molecular Probes) or Streptavidin with Alexa 568 (S11226, Molecular Probes) for 1-2 days at 4° C. Then the brains were washed multiple times with PBS and PBS-T for 3-4 hours and mounted in Vectashield anti-fade mounting media (H-1000, Vector Laboratories). Morphological images were obtained using Nikon A1R MP+ microscope.

### Identification of glomeruli

To identify the glomeruli, we compared the histological images with the atlases provided by Ignell et al., 2005 and Shankar and McMeniman, 2020. Our images matched the Ignell et al., 2005 atlas more closely and hence we have assigned the glomerular identity using this atlas; we have added a prefix ‘I-’ to glomerular names to indicate this. Some glomeruli are comparatively large and invariant in shape and location and could be identified easily in different samples; these served as landmark glomeruli. Some glomeruli were challenging to identify due to unclear boundaries or variations in shape, size, or location. To improve our confidence in the identification, two members from our team independently labeled each histological image. If their initial identifications for an image differed, the two members reanalyzed the image together to arrive at a consensus.

### Behavioral experiments

For behavioral experiments, a custom-built acrylic behavioral chamber was used. It consisted of a main cuboidal chamber (L:60 × B:9 × H:9 cm) and a cylindrical release chamber (L:8 X D:5 cm) that was attached to the center of the rear panel of the main chamber. A rotating door at the junction separated the release chamber from the main chamber. The other end of the release chamber had a copper screen. A sliding double-door was placed in the middle of the main chamber to divide it into two arms on the side and a small middle region between the two doors that connected to the release chamber. Air streams (5L/min) entered from the sides of the chamber through four tubes attached at each side panel. A vacuum tube was placed behind the copper screen of the release chamber to maintain the airflow entering from the sides and exiting at the middle. Odorized air came in from one side and clean air (control) from the other. LED lights were placed at both ends of the main chamber to illuminate both arms equally. During the experiment, a white cardboard cover was placed over the behavioral chamber to avoid visual biases. A black stripe pattern was put on all the sides and bottom of the chamber to provide visual features that might help mosquitoes to navigate. The olfactometer was kept in another acrylic chamber (L:120 × B:60 × H:60 cm) to isolate it from the surrounding. The temperature inside the chamber was maintained between 25°C-30°C and relative humidity between 45%-80%.

For each experiment, 18-20 female mosquitoes were taken from rearing cages and transferred to a starvation cage for 24 hours where they had access to water but not sucrose. Just before the experiments, the mosquitoes were moved to the release chamber and allowed to acclimatize for 30 mins. At the start of the experiment, the odorized stream was switched on and the odor was delivered continuously in pulses of 1 s with gaps of 4 s. After 30 s, the rotating door of the release chamber was opened so that the mosquitoes could fly into the main chamber. Over the next 5 minutes, mosquitoes were allowed to move free in the main chamber. At the end of the experiment, the sliding door was closed and mosquitoes in the two side-arms were counted and considered as responding. The mosquitoes remaining in the release chamber or in the middle part of the main chamber (between the two sliding doors) were also counted and considered to be non-responding. Only those experiments where the number of responding mosquitoes was ≥5 were considered for further analyses. Preference index for the odor was calculated as (N_o_-N_c_)/(N_o_+N_c_), where N_o_ and N_c_ are the numbers of mosquitoes in the odorized arm and the control arm, respectively.

After each experiment, the chamber was cleaned with 70% (v/v) ethanol and flushed with clean air. The side of the odorized air was randomly assigned in each experiment to minimize the bias from any unplanned difference between the left side and the right side of the chamber. The experiments in which experimental conditions or the number of responding mosquitoes were out of the expected range were excluded from further analysis. For each odor, data from 12-14 experiments were obtained.

### Odor delivery

Fourteen odorants were used in this study including eight components of human odor: 1-octen-3-ol (1OCT3OL), 2-pentanone (2PENTONE), 6-methyl-5-hepten-2-one (6MHO), dimethyl trisulfide (DM TS), geranyl acetate (GER ATE), L-lactic acid (L-LA), nonanal (NONANAL), and propionic acid (PR ACID); two plant-derived molecules: (S)-limonene (S-LIMENE) and methyl salicylate (MET SAL); one oviposition attractant (Bentley et al., 1982): 4-methylcyclohexanol (4MCYHOL); one mimicking carbon dioxide (Tauxe et al., 2013): cyclopentanone (CYPTONE); one aggregation pheromone: 4’ethylacetophenone (4EACPONE); and one synthetic repellent: N,N-diethyl-meta-toluamide (DEET). Dilutions (v/v) were done with mineral oil except for L-lactic acid, which was diluted in water. 2 ml of the odorant was placed in a 50-ml glass bottle. For electrophysiology, a stream of dehumidified and filtered compressed air (2L/min) was directed at the animal throughout the recording. The 2L/min flow included a constant stream of 1.8 L/min and a flexible stream of 200 ml/min that passed either through a clean empty vial (during background period) or through the saturated headspace of an odor vial (during odor stimulation), controlled by a 3-way distributor solenoid valve (Product code: 11-13-3-BV-24F88, Parker Hannifin), ensuring that there was no change in the total air flow during odor stimulation. In this arrangement, the odorized stream got diluted by a factor of 0.1 after mixing with constant stream; the dilution indicated against each odor throughout this manuscript is the final dilution coming out of the outlet tube (further dilution caused by mixing of the outlet stream with the ambient air around the animal is not taken into consideration). The outlet was kept 8-10 mm away from the animal. The delivery of odor was confirmed regularly using a photo-ionization detector (200B miniPID, Aurora Scientific). For the behavioral experiments, a constant stream (4.5 L/min) of humidified filtered air was directed into the behavioral chamber from each end of the main chamber. For odor stimulation, an air stream of 0.5 L/min was odorized by passing through an odor vial and added to one side of the chamber, while an equivalent stream of clean air was added on the opposite side. In all the set-ups, odor tubes were replaced frequently to minimize any contamination and odor vials were replaced periodically to ensure stable odorant concentration.

### Morphological Analysis

*Image registration and neuronal similarity analysis:* In order for neuron morphologies acquired from separate brains to be compared, light microscopy images of brains must be registered to a standard template brain space (Bates et al., 2020). For ease of analysis, these neurons can then be ‘skeletonised’ and exported as SWC files. We aimed to register our neurons into an open-source female *Aedes aegypti* brain template (obtained from mosquitobrains.org, courtesy of Meg Younger and Leslie Vosshall). Standard practice in the field is to use the Computational Morphometry ToolKit (CMTK) (Rohlfing and Maurer, 2003) to register brains with a neuropil stain, to a template. However, due to damage to our brains during the recording process, this proved inaccurate. Instead, we performed a landmarks-based registration, marking cross-mapped control points (e.g. specific AL glomeruli, specific tract bends, etc.) between each sample brain and our reference brain, up to 35 control points per brain hemisphere. We used a thin plate spline registration (Bookstein, 1989) implemented in the R package *Morpho* (Schlager, 2017) to create a warping deformation that registered each sample brain to the template, and could be used, via *Morpho* and the *natverse*, to register each of our neuronal reconstructions to the template. Our neuronal reconstructions were of the axonal projections of PNs, and were manually traced using Neutube (Feng et al., 2015) and reconstructed using the SNT plug-in ImageJ. A 3D brain mesh was obtained from the insectbrainDB.org API (Heinze et al., 2021) using the R package *insectbrainr* (Bates et al., 2020). Reconstructed neurons were analysed in R using the *natverse* (Bates et al., 2020). The similarities of the arborizations of the registered PN traces in the protocerebrum neuropil were calculated using NBLAST (Costa et al., 2016). In order to control for overall neuron size, we resampled neurons to a 0.5 micron spacing for each point in the skeleton, and used normalized NBLAST scores (raw NBLAST score for neuron A to B comparison, divided by the score or a self match, i.e. A to A). In order to compare only the morphology of neurons’ axons outside of the mushroom body, we removed all cable from each neuron prior to the most lateral axonal arborization. To remove the influence of cofasciulating tracts on our NBLAST result we also pruned away the higher strahler order cable in these isolated axons, i.e. the central tract, leaving only the axonal branches.

*Glomerular innervation map of LNs*: Since the shape, size and location of glomeruli vary from one sample to another, we focused on 14 landmark glomeruli (I-AD1, I-AD2, I-AM1, I-AM2, I-MD1, I-MD2, I-MD3, I-PC1, I-PD6, I-PL2, I-PL3, I-PL6, I-PM4, and I-V1) whose identity could be confirmed reliably based on one or more of the following features: a specific location, a unique shape, a noticeably larger or smaller size compared to neighboring glomeruli. We manually examined LN arborizations in each of these glomeruli in the image stacks.

### Electrophysiological feature analysis

Spikes were detected from low-pass filtered voltage traces using custom code in MATLAB. Spikes were classified into bursts or isolated spikes: a spike was considered isolated if there were no other spikes within 200 ms on either side of it. Next, for each cell, we extracted following features from the entire duration of the recordings:

i. *Spike amplitude:* For isolated spikes, the distance between the peak of the spike and the left trough or the right trough (whichever gives a larger amplitude) was taken as the amplitude of the spike. For bursts, the amplitude was calculated in the same way for the first and the last spikes of the burst and mean of the two was used. If a cell had isolated spikes, the mean amplitude of isolated spikes was used for further analysis, otherwise the average amplitude from the bursts was used.
ii. *Spike half-width:* The width of the spike at half the height was calculated and averaged over all isolated spikes.
iii. *After-hyperpolarization amplitude:* In band-pass filtered traces (5-500 Hz), we calculated the lowest value of the membrane potential within 10 ms after a spike and from it subtracted the average value of the membrane potential in the window of 20-70 ms after the spike. This value was usually negative and was averaged over all isolated spikes; occasional positive values (due to unexpected fluctuations in the membrane potential) were ignored.
iv. *Isolated spikes fraction:* The number of isolated spikes was divided by the total number of spikes in a cell.

We generated a feature matrix with individual cells as rows and mean values of the above features as columns. Each feature was normalized such that when all the cells are considered the mean of each feature is 0 and the standard deviation is 1. In about 5% of the cases, the feature matrix had missing values (e.g., when a cell did not have any isolated spikes). To enable hierarchical clustering, we interpolated the missing values in the following manner. We made a reference set of cells for which all features were available. For a cell with one or more missing values, we used the available features of that cell to calculate its similarity to each cell in the reference set. Then we estimated a missing feature of that cell as the weighted average of the corresponding values from the reference set, with similarity values as weights.

To include reliable data for this analysis, we manually assigned a quality index on a scale of 0-5 for each cell, based on the resting membrane potential after break-in, background noise (as compared to other cells recorded on same day), average spike size, and the amount of current that had to be injected to stabilize the cell. Cells with quality index greater than 3 were used for this analysis; this corresponded to >80% of morphologically identified neurons (170 of 208 PNs and 42 of 53 LNs).

### Analysis of odor responses

For each cell-odor pair, the response intensity was calculated by subtracting the background firing rate from the firing rate during the response duration for each trial and was then averaged over all trials available for that pair. In each trial the odor valve was turned on at 2 s time point; the first 2 s were used as the background duration and the 2.1-4.1 s interval was used as the response duration (considering 100-150 ms gap between the switching on of the odor valve and the odor actually reaching the animal). Although the odor was delivered for 1 s, we used a 2-s response duration because the odor response often lasted longer than the odor stimulus. Cells that did not show any spike were excluded from this analysis.

For estimating the fractions of cells that responded to an odor or the fraction of odors that activated a given cell, we classified each cell-odor pair as ‘responding’ or ‘non-responding’ using a statistical criterion. We divided the response duration into two 1-s bins and compared the spiking rate in each bin over all trials to the spiking rate in the background period using a signed-rank test. If the test showed a significantly high or low firing rate in either bin compared to the background (P < 0.05), the cell-odor pair was labeled as ‘responding’. The fraction of odors to which a cell responds was determined for only those cells in which at least 5 odors were tested. For PNs, we took the average over all homotypic PNs for each glomerulus. For LNs, we calculated the fractions for individual cells.

### Visualization of the dynamic PN population response as odor trajectories

This analysis was performed using the dataset of 6 common odors that were tested in 64 common PNs. The first 6 s interval of the trial was divided into 60 bins of 100 ms each, and the average firing rate in each bin (minus the background firing rate over the first 2 s) was calculated over all trials for each cell-odor pair. The binned vectors from all 6 odors were concatenated for each PN. Principal component analysis was performed on the resulting 360 × 64 dimensional matrix using the ‘pca’ routine in Matlab. Using the first 3 principal components, we plotted the values for all time bins separately for each odor, yielding 6 odor trajectories in a 3-D space.

### Odor classification based on PN responses

We performed a classification accuracy analysis (Stopfer et al., 2003; Gupta and Stopfer, 2014) using the spiking responses of the PN-odor set (64 PNs × 6 odors). As this analysis used the temporal patterns of responses, we ignored the first trial for each cell-odor pair because for some odors the temporal profile of odor delivery was slightly different in the first trial compared to other trials; we used the next 6 trials for each cell-odor pair. For each trial, the spikes in the response duration (2.1-4.1 s) were divided into 8 bins of 250 ms each and the firing rate in each bin was calculated after subtracting the background firing rate. For each odor, we generated a matrix with 6 rows (corresponding to 6 trials) and 64 × 8 columns, corresponding to 8 bins from 64 cells. The first 64 columns included the values from the first bins of all 64 cells, the next 64 columns from the second bins, and so on. We estimated classification accuracy for different response durations (i.e., for different numbers of bins used). One trial was selected as test data and the remaining trials were used as training data. To calculate accuracy for a given number of bins (say *k*), we generated a template for each odor by taking the centroid of the training trials using the first *k* bins. Next, we calculated Euclidean distances between the test data of one odor (using the first *k* bins) and the templates of all odors. If the test data was closest to the template of the same odor, the accuracy was taken as 1, otherwise 0. The accuracy was calculated for all test odors, and then averaged. This was repeated six times by using a different trial as the test trial each time and the mean accuracy was calculated. We then calculated the cumulative accuracy for different response durations by varying *k* from 1 to 8.

We also calculated the classification accuracy as a function of the number of cells while using the first 3 bins (750 ms response duration). We varied the number of cells (*n*) from 5 to 60 in gaps of 5. For each value of *n*, we performed the classification analysis 10 times, each time sampling a random set of *n* cells, and then took the average accuracy value over the 10 iterations.

### Behavior-PN correlations

For each pair of odors, we calculated the difference in their preference indices (ΔPI). We generated a response vector for each odor in the pair by concatenating the responses from PNs that were recorded for both the odors (again, the response was quantified as the firing rate in the 2.1-4.1 s response duration minus the background firing rate, averaged over all trials). Then, we calculated the Pearson correlation coefficient (*R*) between the two odor vectors and used 1 - R as the distance between the PN responses for the odor pair. Next, the PN-behavior correlation was calculated as the Pearson correlation between the ΔPI and 1 - *R* values over all odor pairs. We also calculated the PN-behavior correlation as a function of the number of common PNs (*n*). For a given *n*, we randomly selected *n* PNs from the common PNs for each odor pair and calculated the PN-behavior correlation from this reduced dataset (this random sampling was repeated 20 times for each *n* and the average correlation was obtained). For large values of *n*, some odor pairs had fewer than *n* common PNs and were ignored in the calculations.

## Data availability

The morphological reconstructions of PNs registered to the template brain are available at https://github.com/neuralsystems/MosquitoAL. All other datasets are available from the corresponding author upon request.

## Code availability

Code used for analyses are available from the corresponding author on request. The classification analysis was performed using a custom library available at https://github.com/neuralsystems/temporal_classification.

## Acknowledgements

We thank Arun Shankar for maintaining the mosquito facility, Anubhav Kumar and Ranjeet Kumar for aiding in behavioral experiments, and Kazumichi Shimizu for advice on electrophysiology and histology. We thank Meg Younger and Leslie Vosshall for sharing the female *Aedes aegypti* brain template on the mosquitobrains.org portal. This work was supported by the DBT/Wellcome Trust India Alliance Fellowship [grant number IA/I/15/2/502091], DST/SERB Swarnajayanti Fellowship [SB/SJF/2021-22/04-C], and SERB Core Research Grant [CRG/2020/004719] awarded to N.G.

## Authors Contributions

Conceptualization: NG and PS Methodology: PS, ASB and NG

Data collection (Electrophysiological): PS and Shefali G Data collection (Behavioral): PS, Sanket G and VR

Data collection (Morphological): PS, Shefali G and AKG Data curation: PS and Shefali G

Data analysis: PS, Smith G, AT, ASB, Sanket G and NG Writing: PS, NG and ASB

## References

1. Afify A, Potter CJ (2020) Insect repellents mediate species-specific olfactory behaviours in mosquitoes. Malar J 19:127.

2. Anton S, Van Loon JJA, Meijerink J, Smid HM, Takken W, Rospars JP (2003) Central projections of olfactory receptor neurons from single antennal and palpal sensilla in mosquitoes. Arthropod Struct Dev 32:319–327.

3. Aubry F et al. (2020) Enhanced Zika virus susceptibility of globally invasive Aedes aegypti populations. Science (80-) 370:991–996.

4. Bates AS, Schlegel P, Roberts RJV, Drummond N, Tamimi IFM, Turnbull R, Zhao X, Marin EC, Popovici PD, Dhawan S, Jamasb A, Javier A, Serratosa Capdevila L, Li F, Rubin GM, Waddell S, Bock DD, Costa M, Jefferis GSXE (2020) Complete Connectomic Reconstruction of Olfactory Projection Neurons in the Fly Brain. Curr Biol 30:3183–3199.e6.

5. Bentley MD, McDaniel IN, Davis EE (1982) Studies of 4-methylcyclohexanol: An Aedes triseriatus (Diptera: Culicidae) oviposition attractant. J Med Entomol 19:589–592.

6. Bernier UR, Booth MM, Yost RA (1999) Analysis of human skin emanations by gas chromatography/mass spectrometry. 1. Thermal desorption of attractants for the yellow fever mosquito (aedes aegypti) from handled glass beads. Anal Chem 71:1–7.

7. Bernier UR, Kline DL, Allan SA, Barnard DR (2007) Laboratory Comparison of Aedes Aegypti Attraction to Human Odors and to Synthetic Human Odor Compounds and Blends. J Am Mosq Control Assoc 23:288–293.

8. Bernier UR, Kline DL, Barnard DR, Schreck CE, Yost RA (2000) Analysis of human skin emanations by gas chromatography/mass spectrometry. 2. Identification of volatile compounds that are candidate attractants for the yellow fever mosquito (Aedes aegypti). Anal Chem 72:747–756.

9. Bisch-Knaden S, Dahake A, Sachse S, Knaden M, Hansson BS (2018) Spatial Representation of Feeding and Oviposition Odors in the Brain of a Hawkmoth. Cell Rep 22:2482–2492.

10. Bohbot JD, Sparks JT, Dickens JC (2014) The maxillary palp of Aedes aegypti, a model of multisensory integration. Insect Biochem Mol Biol 48:29–39.

11. Bookstein FL (1989) Principal Warps: Thin-Plate Splines and the Decomposition of Deformations. IEEE Trans Pattern Anal Mach Intell 11:567–585.

12. Cardé RT (2015) Multi-Cue Integration: How Female Mosquitoes Locate a Human Host. Curr Biol 25:R793–R795.

13. Carey AF, Wang G, Su CY, Zwiebel LJ, Carlson JR (2010) Odorant reception in the malaria mosquito Anopheles gambiae. Nature 464:66–71.

14. Choi GB, Stettler DD, Kallman BR, Bhaskar ST, Fleischmann A, Axel R (2011) Driving opposing behaviors with ensembles of piriform neurons. Cell 146:1004–1015.

15. Chou Y-H, Spletter ML, Yaksi E, Leong JCS, Wilson RI, Luo L (2010) Diversity and wiring variability of olfactory local interneurons in the Drosophila antennal lobe. Nat Neurosci 13:439–449.

16. Cork A, Park KC (1996) Identification of electrophysiologically-active compounds for the malaria mosquito, Anopheles gambiae, in human sweat extracts. Med Vet Entomol 10:269–276.

17. Costa M, Manton JD, Ostrovsky AD, Prohaska S, Jefferis GSXE (2016) NBLAST: Rapid, Sensitive Comparison of Neuronal Structure and Construction of Neuron Family Databases. Neuron 91:293–311.

18. Couto A, Alenius M, Dickson BJ (2005) Molecular, anatomical, and functional organization of the Drosophila olfactory system. Curr Biol 15:1535–1547.

19. Das A, Sen S, Lichtneckert R, Okada R, Ito K, Rodrigues V, Reichert H (2008) Drosophila olfactory local interneurons and projection neurons derive from a common neuroblast lineage specified by the empty spiracles gene. Neural Dev 3:1–17.

20. Davis EE, Sokolove PG (1976) Lactic acid-sensitive receptors on the antennae of the mosquito, Aedes aegypti. J Comp Physiol 105:43–54.

21. Distler PG, Boeckh J (1997) Central projections of the maxillary and antennal nerves in the mosquito Aedes aegypti. J Exp Biol 200:1873–1879.

22. Dormont L, Bessière JM, Cohuet A (2013) Human Skin Volatiles: A Review. J Chem Ecol 39:569–578.

23. Dweck HKM, Ebrahim SAM, Thoma M, Mohamed AAM, Keesey IW, Trona F, Lavista-Llanos S, Svatoš A, Sachse S, Knaden M, Hansson BS (2015) Pheromones mediating copulation and attraction in Drosophila. Proc Natl Acad Sci U S A 112:E2829–E2835.

24. Fawaz EY, Allan SA, Bernier UR, Obenauer PJ, Diclaro JW (2014) Swarming mechanisms in the yellow fever mosquito: aggregation pheromones are involved in the mating behavior of Aedes aegypti. J Vector Ecol 39:347–354.

25. Frechter S, Bates AS, Tootoonian S, Dolan MJ, Manton J, Jamasb AR, Kohl J, Bock D, Jefferis G (2019) Functional and anatomical specificity in a higher olfactory centre. Elife 8.

26. Fusca D, Kloppenburg P (2021) Task-specific roles of local interneurons for inter- and intraglomerular signaling in the insect antennal lobe. Elife 10:1–19.

27. Gallagher M, Wysocki CJ, Leyden JJ, Spielman AI, Sun X, Preti G (2008) Analyses of volatile organic compounds from human skin. Br J Dermatol 159:780–791.

28. Geier M, Bosch OJ, Boeckh J (1999) Ammonia as an attractive component of host odour for the yellow fever mosquito, Aedes aegypti. Chem Senses 24:647–653.

29. Ghaninia M, Hansson BS, Ignell R (2007a) The antennal lobe of the African malaria mosquito, Anopheles gambiae -innervation and three-dimensional reconstruction. Arthropod Struct Dev 36:23–39.

30. Ghaninia M, Ignell R, Hansson BS (2007b) Functional classification and central nervous projections of olfactory receptor neurons housed in antennal trichoid sensilla of female yellow fever mosquitoes, Aedes aegypti. Eur J Neurosci 26:1611–1623.

31. Ghaninia M, Larsson M, Hansson BS, Ignell R (2008) Natural odor ligands for olfactory receptor neurons of the female mosquito Aedes aegypti: use of gas chromatography-linked single sensillum recordings. J Exp Biol 211:3020–3027.

32. Gupta N, Stopfer M (2014) A temporal channel for information in sparse sensory coding. Curr Biol 24:2247–2256.

33. Hallem EA, Carlson JR (2006) Coding of Odors by a Receptor Repertoire. Cell 125:143–160.

34. Haverkamp A, Hansson BS, Knaden M (2018) Combinatorial codes and labeled lines: How insects use olfactory cues to find and judge food, mates, and oviposition sites in complex environments. Front Physiol 9:1–8.

35. Heinze S, El Jundi B, Berg BG, Homberg U, Menzel R, Pfeiffer K, Hensgen R, Zittrell F, Dacke M, Warrant E, Pfuhl G, Rybak J, Tedore K (2021) A unified platform to manage, share, and archive morphological and functional data in insect neuroscience. Elife 10:1– 25.

36. Hill SR, Hansson BS, Ignell R (2009) Characterization of antennal trichoid sensilla from female Southern house mosquito, culex quinquefasciatus say. Chem Senses 34:231–252.

37. Homberg U, Montague RA, Hildebrand JG (1988) Anatomy of antenno-cerebral pathways in the brain of the sphinx moth Manduca sexta. Cell Tissue Res 254:255–281.

38. Huang J, Zhang W, Qiao W, Hu A, Wang Z (2010) Functional connectivity and selective odor responses of excitatory local interneurons in drosophila antennal lobe. Neuron 67:1021– 1033.

39. Huoviala P, Dolan M-J, Love F, Frechter S, Roberts RJ V, Mitrevica Z, Schlegel P, Bates ASS, Aso Y, Rodrigues T, Cornwall H, Stensmyr M, Bock D, Rubin GM, Costa M, Jefferis GSXE (2020) Neural circuit basis of aversive odour processing inDrosophilafrom sensory input to descending output. bioRxiv:394403.

40. Ignell R, Dekker T, Ghaninia M, Hansson BS (2005) Neuronal architecture of the mosquito deutocerebrum. J Comp Neurol 493:207–240.

41. Kuebler LS, Olsson SB, Weniger R, Hansson BS (2011) Neuronal processing of complex mixtures establishes a unique odor representation in the moth antennal lobe. Front Neural Circuits 5:7.

42. Kurtovic A, Widmer A, Dickson BJ (2007) A single class of olfactory neurons mediates behavioural responses to a Drosophila sex pheromone. Nature 446:542–546.

43. Kymre JH, Berge CN, Chu X, Ian E, Berg BG (2021) Antennal-lobe neurons in the moth Helicoverpa armigera: Morphological features of projection neurons, local interneurons, and centrifugal neurons. J Comp Neurol 529:1516–1540.

44. Lahondère C, Vinauger C, Okubo RP, Wolff GH, Chan JK, Akbari OS, Riffell JA (2020) The olfactory basis of orchid pollination by mosquitoes. Proc Natl Acad Sci U S A 117:708– 716.

45. Lai SL, Awasaki T, Ito K, Lee T (2008) Clonal analysis of Drosophila antennal lobe neurons: Diverse neuronal architectures in the lateral neuroblast lineage. Development 135:2883– 2893.

46. Leal WS (2014) The enigmatic reception of DEET - The gold standard of insect repellents. Curr Opin Insect Sci 6:93–98.

47. Lei H, Reisenman CE, Wilson CH, Gabbur P, Hildebrand JG (2011) Spiking patterns and their functional implications in the antennal lobe of the tobacco hornworm Manduca sexta. PLoS One 6.

48. Liu MZ, Vosshall LB (2019) General Visual and Contingent Thermal Cues Interact to Elicit Attraction in Female Aedes aegypti Mosquitoes. Curr Biol 29:2250–2257.e4.

49. Liu WW, Wilson RI (2013) Glutamate is an inhibitory neurotransmitter in the Drosophila olfactory system. Proc Natl Acad Sci U S A 110:10294–10299.

50. Logan JG, Birkett MA, Clark SJ, Powers S, Seal NJ, Wadhams LJ, Mordue AJ, Pickett JA (2008) Identification of human-derived volatile chemicals that interfere with attraction of Aedes aegypti mosquitoes. J Chem Ecol 34:308–322.

51. Logan JG, Stanczyk NM, Hassanali A, Kemei J, Santana AEG, Ribeiro KAL, Pickett JA, Mordue AJ (2010) Arm-in-cage testing of natural human-derived mosquito repellents. Malar J 9:239.

52. Lu T, Qiu YT, Wang G, Kwon JY, Rutzler M, Kwon HW, Pitts RJ, van Loon JJ a, Takken W, Carlson JR, Zwiebel LJ (2007) Odor Coding in the Maxillary Palp of the Malaria Vector Mosquito Anopheles gambiae. Curr Biol 17:1533–1544.

53. Majeed S, Hill SR, Birgersson G, Ignell R (2016) Detection and perception of generic host volatiles by mosquitoes modulate host preference: Context dependence of (R)-1-octen-3-ol. R Soc Open Sci 3.

54. Malnic B, Hirono J, Sato T, Buck LB (1999) Combinatorial receptor codes for odors. Cell 96:713–723.

55. Masse NY, Turner GC, Jefferis GSXE (2009) Olfactory Information Processing in Drosophila. Curr Biol 19:R700–R713.

56. McBride CS, Baier F, Omondi AB, Spitzer SA, Lutomiah J, Sang R, Ignell R, Vosshall LB (2014) Evolution of mosquito preference for humans linked to an odorant receptor. Nature 515:222–227.

57. McMeniman CJ, Corfas RA, Matthews BJ, Ritchie SA, Vosshall LB (2014) Multimodal integration of carbon dioxide and other sensory cues drives mosquito attraction to humans. Cell 156:1060–1071.

58. Melo N, Wolff GH, Costa-da-Silva AL, Arribas R, Triana MF, Gugger M, Riffell JA, DeGennaro M, Stensmyr MC (2020) Geosmin Attracts Aedes aegypti Mosquitoes to Oviposition Sites. Curr Biol 30:127–134.e5.

59. Meyer A, Giovanni Galizia C, Nawrot MP (2013) Local interneurons and projection neurons in the antennal lobe from a spiking point of view. J Neurophysiol 110:2465–2474.

60. Mobbs PG (1982) The brain of the honeybee Apis mellifera. I. The connections and spatial organization of the mushroom bodies. Philos Trans R Soc London 298:309–354.

61. Mohamed AAM, Retzke T, Das Chakraborty S, Fabian B, Hansson BS, Knaden M, Sachse S (2019) Odor mixtures of opposing valence unveil inter-glomerular crosstalk in the Drosophila antennal lobe. Nat Commun 10:1–17.

62. Mozūraitis R, Hajkazemian M, Zawada JW, Szymczak J, Pålsson K, Sekar V, Biryukova I, Friedländer MR, Koekemoer LL, Baird JK, Borg-Karlson AK, Emami SN (2020) Male swarming aggregation pheromones increase female attraction and mating success among multiple African malaria vector mosquito species. Nat Ecol Evol 4:1395–1401.

63. Obaldia ME De, Morita T, Dedmon LC, Boehmler DJ, Jiang CS, Zeledon E V., Cross JR, Vosshall LB (2022) Differential mosquito attraction to humans is associated with skin-derived carboxylic acid levels. bioRxiv:2022.01.05.475088.

64. Peach DAH, Gries R, Young N, Lakes R, Galloway E, Alamsetti SK, Ko E, Ly A, Gries G (2019) Attraction of female Aedes Aegypti (L.) to aphid honeydew. Insects 10.

65. Ponnusamy L, Xu N, Nojima S, Wesson DM, Schal C, Apperson CS (2008) Identification of bacteria and bacteria-associated chemical cues that mediate oviposition site preferences by Aedes aegypti. Proc Natl Acad Sci U S A 105:9262–9267.

66. Raji JI, Melo N, Castillo JS, Gonzalez S, Saldana V, Stensmyr MC, DeGennaro M (2019) Aedes aegypti Mosquitoes Detect Acidic Volatiles Found in Human Odor Using the IR8a Pathway. Curr Biol 29:1253–1262.e7.

67. Raman B, Joseph J, Tang J, Stopfer M (2010) Temporally diverse firing patterns in olfactory receptor neurons underlie spatiotemporal neural codes for odors. J Neurosci 30:1994–2006.

68. Riabinina O, Task D, Marr E, Lin CC, Alford R, O’Brochta DA, Potter CJ (2016) Organization of olfactory centres in the malaria mosquito Anopheles gambiae. Nat Commun 7:1–12.

69. Rohlfing T, Maurer CR (2003) Nonrigid image registration in shared-memory multiprocessor environments with application to brains, breasts, and bees. IEEE Trans Inf Technol Biomed 7:16–25.

70. Rudolphs W (1922) Chemotropism of mosquitoes. Bull NJ Agric Exp Stn 367:4–23.

71. Ruta V, Datta SR, Vasconcelos ML, Freeland J, Looger LL, Axel R (2010) A dimorphic pheromone circuit in Drosophila from sensory input to descending output. Nature 468:686–690.

72. Saha D, Raman B (2015) Relating early olfactory processing with behavior: A perspective. Curr Opin Insect Sci 12:54–63.

73. Schlager S (2017) Morpho and Rvcg - Shape Analysis in R: R-Packages for Geometric Morphometrics, Shape Analysis and Surface Manipulations, 1st ed. Elsevier Ltd.

74. Schlegel P, Bates AS, Stürner T, Jagannathan SR, Drummond N, Hsu J, Capdevila LS, Javier A, Marin EC, Barth-Maron A, Tamimi IFM, Li F, Rubin GM, Plaza SM, Costa M, Jefferis GSXE (2021) Information flow, cell types and stereotypy in a full olfactory connectome. Elife 10:1–47.

75. Seki Y, Kanzaki R (2008) Comprehensive morphological identification and GABA immunocytochemistry of antennal lobe local interneurons in Bombyx mori. J Comp Neurol 506:93–107.

76. Shang Y, Claridge-Chang A, Sjulson L, Pypaert M, Miesenböck G (2007) Excitatory Local Circuits and Their Implications for Olfactory Processing in the Fly Antennal Lobe. Cell 128:601–612.

77. Shankar S, McMeniman CJ (2020) An updated antennal lobe atlas for the yellow fever mosquito aedes aegypti.

78. Shankar S, Tauxe GM, Spikol ED, Li M, Akbari OS, Giraldo D, Mcmeniman CJ (2020) Synergistic Coding of Human Odorants in the Mosquito Brain. bioRxiv:2–43.

79. Singh SS, Mittal AM, Chepurwar S (2019) Olfactory Concepts of Insect Control - Alternative to insecticides. Olfactory Concepts Insect Control - Altern to Insectic.

80. Smith CN, Smith N, Gouck HK, Weidhaas DE, Gilbert IH, Mayer MS, Smittle BJ, Hofbauer A (1970) L-lactic acid as a factor in the attraction of Aedes aegypti (Diptera: Culicidae) to human hosts. Ann Entomol Soc Am 63:760–770.

81. Stensmyr MC, Dweck HKM, Farhan A, Ibba I, Strutz A, Mukunda L, Linz J, Grabe V, Steck K, Lavista-Llanos S, Wicher D, Sachse S, Knaden M, Becher PG, Seki Y, Hansson BS (2012) A conserved dedicated olfactory circuit for detecting harmful microbes in drosophila. Cell 151:1345–1357.

82. Stocker RF, Lienhard MC, Borst A, Fischbach KF (1990) Neuronal architecture of the antennal lobe in Drosophila melanogaster. Cell Tissue Res 262:9–34.

83. Stopfer M, Jayaraman V, Laurent G (2003) Intensity versus identity coding in an olfactory system. Neuron 39:991–1004.

84. Syed Z, Leal WS (2009) Acute olfactory response of Culex mosquitoes to a human- and bird-derived attractant. Proc Natl Acad Sci U S A 106:18803–18808.

85. Taisz I, Donà E, Münch D, Bailey SN, Morris WJ, Meechan KI, Stevens KM, Varela I, Gkantia M, Schlegel P, Ribeiro C, Jefferis GSXE, Galili DS (2022) Generating parallel representations of position and identity in the olfactory system. bioRxiv 13:2022.05.13.491877.

86. Tauxe GM, Macwilliam D, Boyle SM, Guda T, Ray A (2013) Targeting a dual detector of skin and CO2 to modify mosquito host seeking. Cell 155:1365–1379.

87. Van Breugel F, Riffell J, Fairhall A, Dickinson MH (2015) Mosquitoes use vision to associate odor plumes with thermal targets. Curr Biol 25:2123–2129.

88. Verhulst NO, Umanets A, Weldegergis BT, Maas JPA, Visser TM, Dicke M, Smidt H, Takken W (2018) Do apes smell like humans? The role of skin bacteria and volatiles of primates in mosquito host selection. J Exp Biol 221.

89. Vinauger C, Lahondère C, Wolff GH, Locke LT, Liaw JE, Parrish JZ, Akbari OS, Dickinson MH, Riffell JA (2018) Modulation of Host Learning in Aedes aegypti Mosquitoes. Curr Biol 28:333–344.e8.

90. Vinauger C, Van Breugel F, Locke LT, Tobin KKS, Dickinson MH, Fairhall AL, Akbari OS, Riffell JA (2019) Visual-Olfactory Integration in the Human Disease Vector Mosquito Aedes aegypti. Curr Biol 29:2509–2516.e5.

91. Warth Pérez Arias CC, Frosch P, Fiala A, Riemensperger TD (2020) Stochastic and Arbitrarily Generated Input Patterns to the Mushroom Bodies Can Serve as Conditioned Stimuli in Drosophila. Front Physiol 11:53.

92. Wilson RI, Turner GC, Laurent G (2004) Transformation of Olfactory Representations in the Drosophila Antennal Lobe. Science (80-) 303:366–371.

93. Wong AM, Wang JW, Axel R (2002) Spatial representation of the glomerular map in the Drosophila protocerebrum. Cell 109:229–241.

94. Ye Z, Liu F, Liu N (2021a) Three-dimensional structure of the antennal lobe in the Southern house mosquito Culex quinquefasciatus. Insect Sci 28:93–102.

95. Ye Z, Liu F, Sun H, Baker A, Zwiebel LJ (2021b) Discrete Roles of the Ir76b Ionotropic Co-Receptor Impact Olfaction, Blood Feeding, and Mating in the Malaria Vector Mosquito Anopheles coluzzii. bioRxiv 119:2021.07.05.451160.

96. Zhao Z, Zung JL, Hinze A, Kriete AL, Iqbal A, Younger MA, Matthews BJ, Merhof D, Thiberge S, Ignell R, Strauch M, McBride CS (2022) Mosquito brains encode unique features of human odour to drive host seeking. Nature 605:706–712.

