## Supplementary Information for "Combinatorial encoding of odors in the mosquito antennal lobe"

Singh et al.

**Supplementary Figure 1**

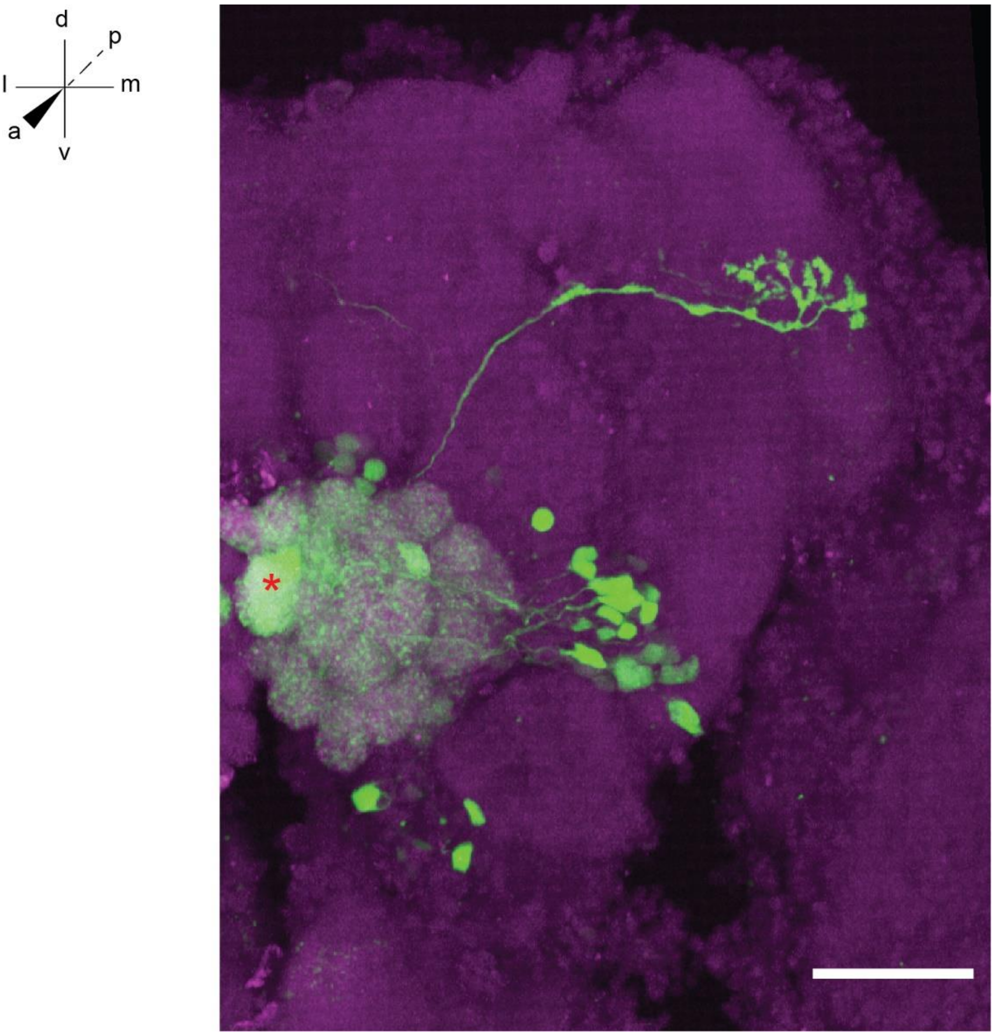


**Supplementary Figure 1: Untargeted labeling of cell bodies**

The maximum intensity projection from an image stack of one half of an adult female *Aedes aegypti* brain showing the antennal lobe and higher brain regions. One PN was recorded and filled on this side of the brain (corresponding to the brightly filled glomerulus indicated by asterisk and the axonal projection to the higher brain areas). Faint signals in the entire antennal lobe and in some cell bodies in the lateral and ventral clusters are also observed, suggesting the presence of gap junctions between the recorded PN and other LNs/PNs. Co-ordinate axes: dorsal-ventral (d-v), anterior-posterior (a-p), and medial-lateral (m-l). Green: biocytin; magenta: Dncad (neuropil marker). Scale bar, 50 µm.

### Supplementary Figure 2

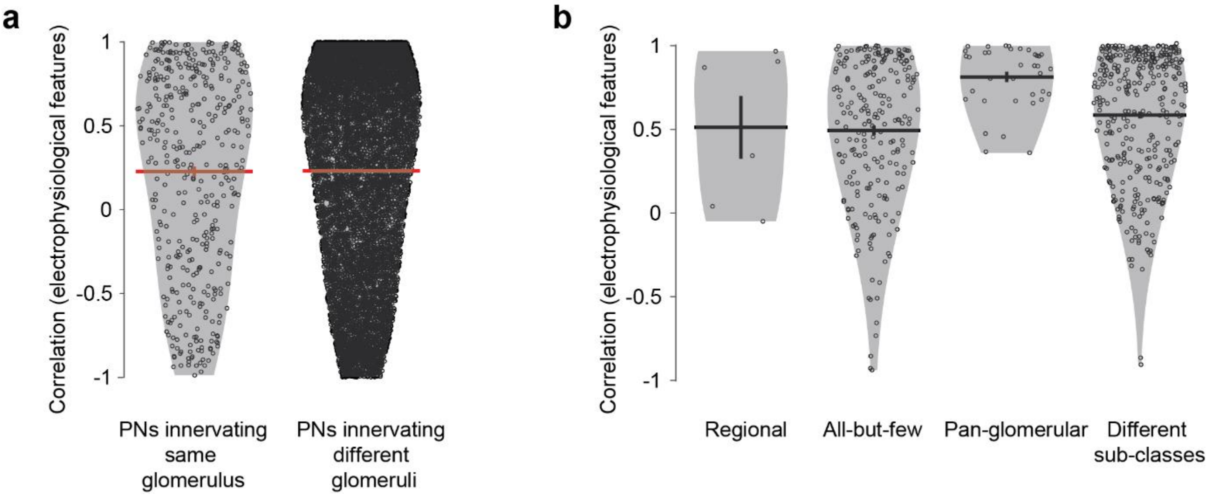


**Supplementary Figure 2: Electrophysiological characterization of sub-classes of PNs and LNs**

**a** Correlations between the electrophysiological features of pairs of PNs innervating the same glomerulus (n = 418) or innervating different glomeruli (n = 10313). **b** Correlations between the electrophysiological features of pairs of LNs within *regional* (n = 6), *all-but-few* (n = 231), or *pan-glomerular* (n = 36) sub-classes, or pairs of LNs belonging to different sub-classes (n = 357). Error bar represents s.e.m.

**Supplementary Figure 3**

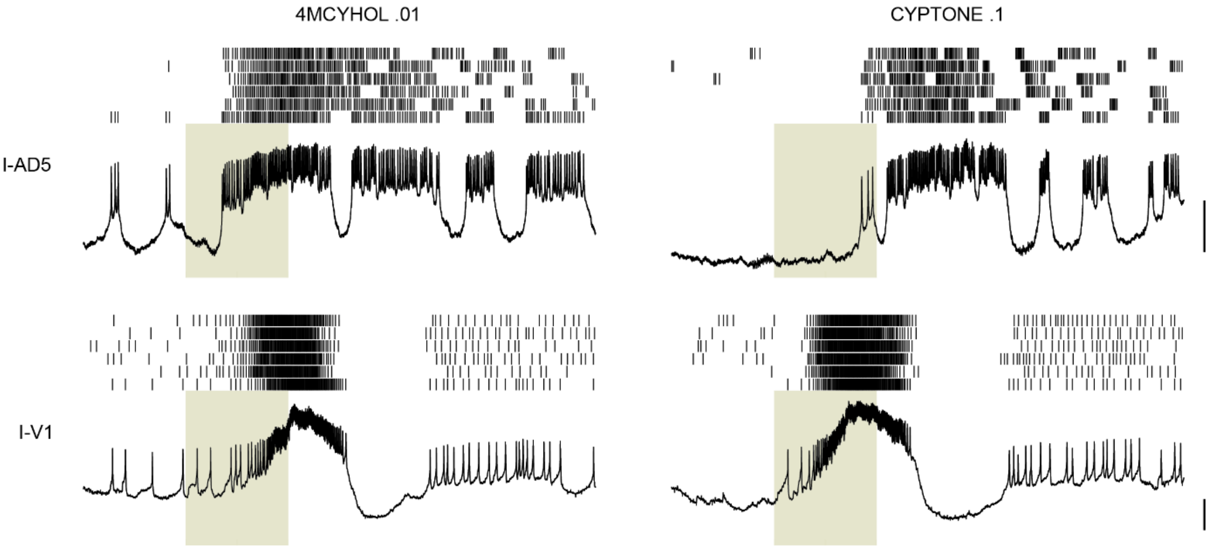


**Supplementary Figure 3: Delayed onset of odor response**

Example of PNs showing a delayed onset of odor responses. Individual panels show 5s-long recording snippets (including the trace from the first trial and spike raster for all trials) obtained from two PNs tested with two odors. The onset of spiking in I-AD5 PN in response to CYPTONE .1 is delayed by ~800 ms after the start of the odor delivery while the response of the same PN to 4MCYHOL .01 starts much earlier. In contrast, in the I-V1 PN, the response to CYPTONE .1 starts earlier than the response to 4MCYHOL .01**.** Scale bars, 10 mV.

**Supplementary Figure 4**

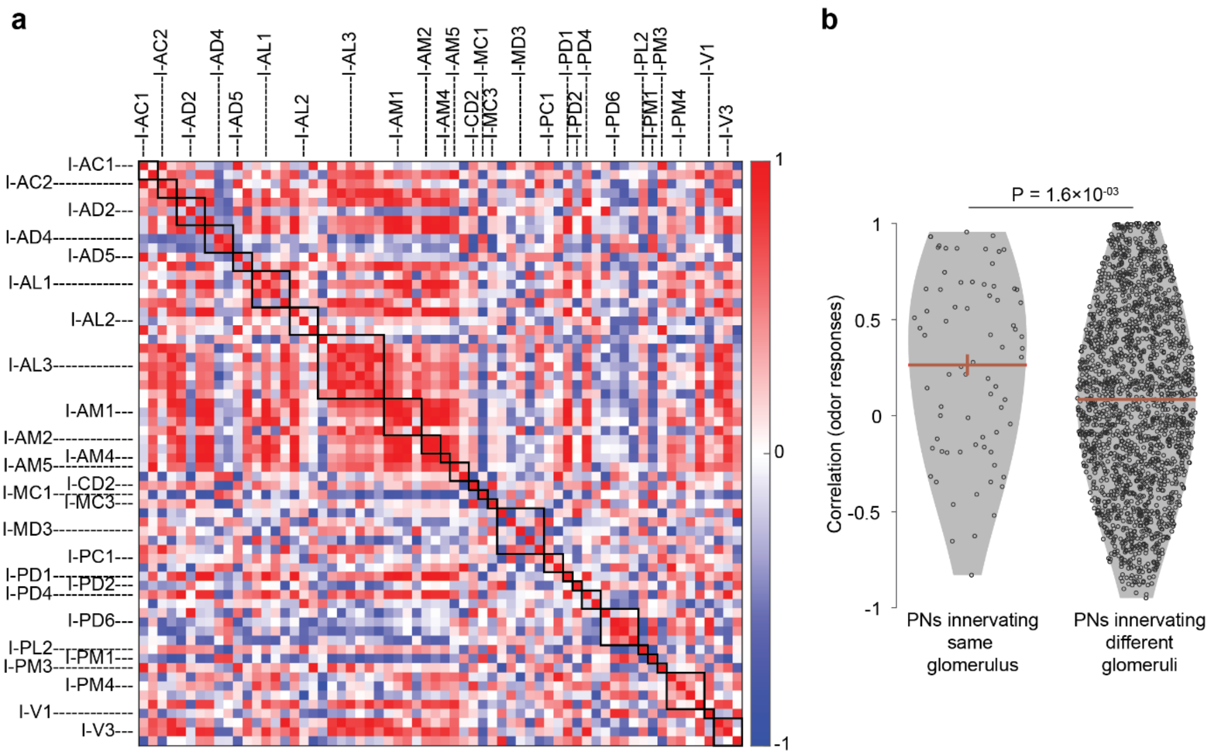


**Supplementary Figure 4: Odor response similarity between homotypic PNs**

**a** Pairwise correlations between vectors of odor responses of PNs. Responses of 64 PNs to 6 frequently tested odors were used. PNs are arranged according to glomerular identity; boxes along the diagonal indicate PNs belonging to the same glomerulus. **b** Comparison of the correlation values from **(a)** for PN pairs innervating the same glomerulus (left) (n = 74) or innervating different glomeruli (right) (n = 1942).

**Supplementary Figure 5**

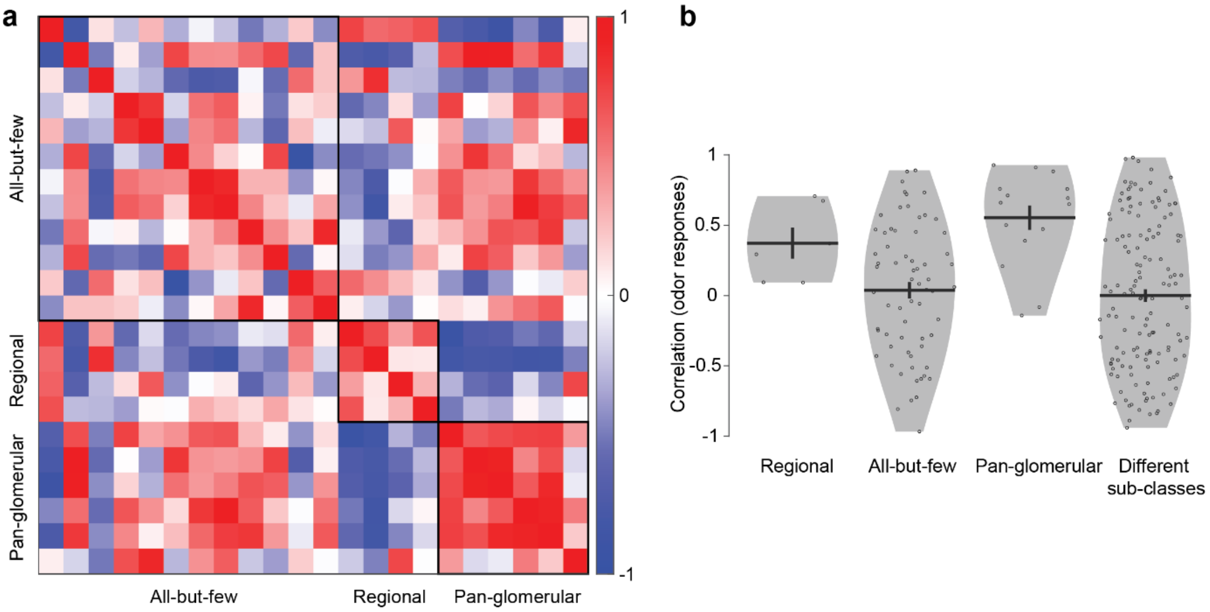


**Supplementary Figure 5: Odor response similarity between morphological sub-classes of LNs**

**a** Pairwise correlations between vectors of odor responses of LNs. Responses of 22 LNs to 6 frequently tested odors were used. Boxes indicate morphological sub-classes of LNs. **b** Comparison of the correlation values from **(a)** for pairs of LNs within *regional* (n = 6), *all-but-few* (n = 66), or *pan-glomerular* (n = 15) sub-classes, or pairs of LNs belonging to different sub-classes (n = 144).

### Supplementary Figure 6

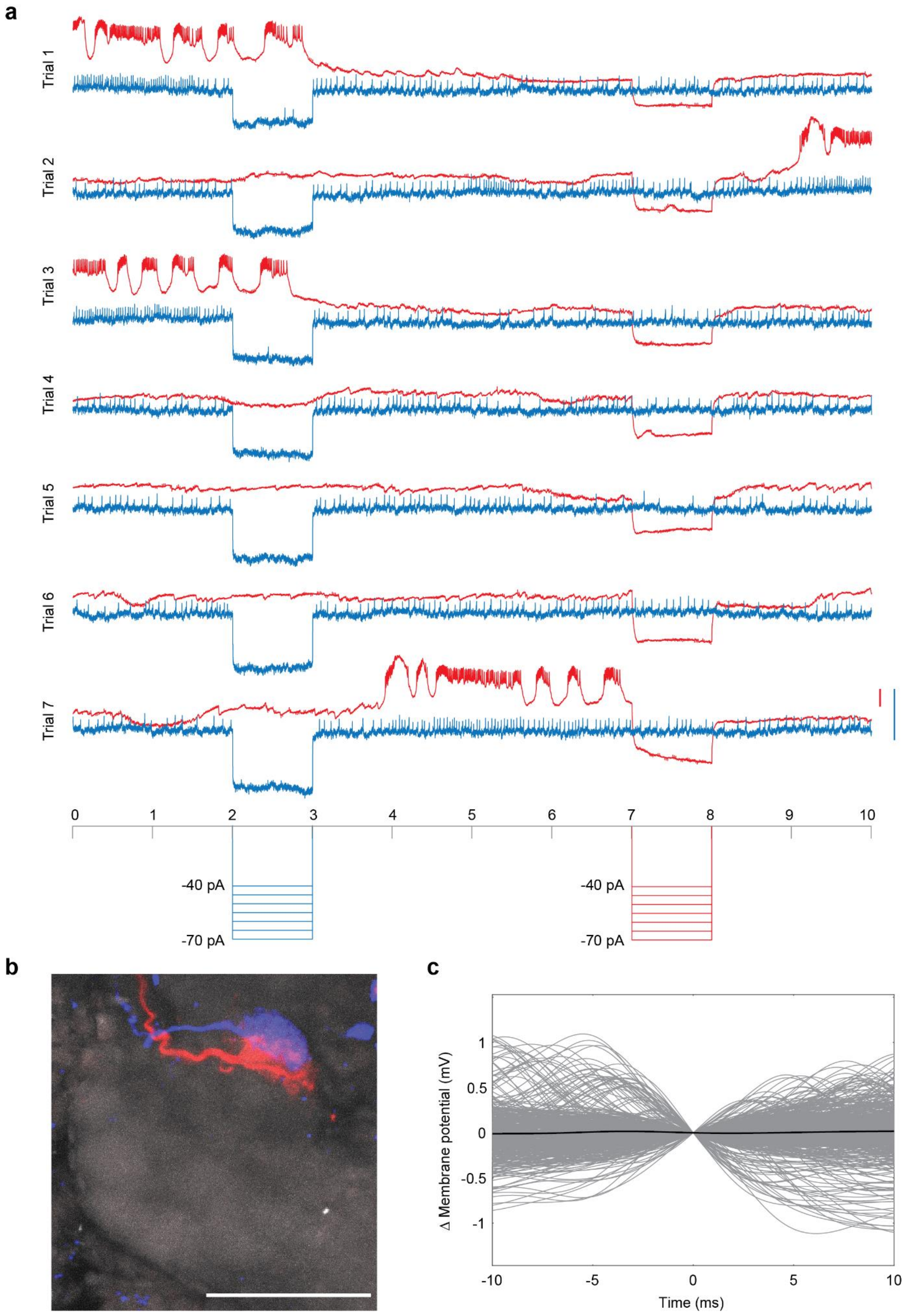

**Supplementary Figure 6: Indirect lateral interaction between I-AL3 and I-PL2 PNs**

**a** Simultaneous recording from an I-AL3 (blue) and an I-PL2 (red) PN. Pulses of hyperpolarizing current (-40 pA in the first trial, increasing in steps of -5 pA in every trial) were injected during 2-3 s in I-AL3 (blue) and during 7-8 s in I-PL2 (red). No effect is seen in either cell when the other was hyperpolarized. **b** Maximum intensity projection of an image stack of the antennal lobe showing the glomerular innervations of the two PNs recorded simultaneously. Note that the two glomeruli are adjacent to each other (they appear overlapping in the flattened stack but are non-overlapping in 3-D). Blue: I-AL3, red: I-PL2, grey: Dncad (neuropil marker). Scale bar, 50 µm. **c** Spike-triggered average of the I-AL3 PN's membrane potential, triggered on the I-PL2 PN spikes, did not provide any evidence of a direct connection from the I-PL2 PN to the I-AL3 PN. Grey: individual traces; black: average of all traces.
